# Cryo-EM structures of naturally occurring dimeric photosystem II complexes lacking the Mn_4_CaO_5_ cluster

**DOI:** 10.1101/2025.07.17.665390

**Authors:** Ziyu Zhao, Irene Vercellino, Julian P. Whitelegge, Karim Maghlaoui, Peter J. Nixon, Leonid A. Sazanov

## Abstract

Robust oxygenic photosynthesis requires the efficient assembly and repair of the multi-subunit oxygen-evolving photosystem II (PSII) complex. Previous electron cryo-microscopy (cryo-EM) structures of PSII assembly/disassembly intermediates have relied on the analysis of deletion mutants or removal of PSII subunits *in vitro*. Here we have determined the cryo-EM structures of naturally occurring dimeric PSII intermediates from the cyanobacterium *Thermosynechococcus vestitus* at a resolution of 2.2 Å. These intermediates contain inactive dimers lacking the oxygen-evolving complex (OEC) and semi-active dimers with the OEC present in one of the two monomers. Our structural data provide a mechanism for how assembly and disassembly of the Mn_4_CaO_5_ cluster is coordinated with the binding and release of the extrinsic proteins: restructuring of the C-terminal tail of D1 subunit during assembly or disassembly of the Mn cluster triggers conformational changes in D2, CP47 and CP43 to drive the binding/release of the extrinsic proteins. A combination of structural and mass spectrometry data indicates that the inactive PSII complexes are disassembly complexes and that oxidation of D1-His332, a monodentate ligand to one of the Mn ions of the OEC, is an early event in the photoinactivation of PSII.

## INTRODUCTION

Photosystem II (PSII) is the light-driven multi-subunit membrane complex found in cyanobacteria and chloroplasts that catalyzes the oxidation of water and reduction of plastoquinone during oxygenic photosynthetic electron transport. PSII is the only enzyme known in nature that is able to catalyze the water-splitting reaction, making it essential for producing oxygen and supporting aerobic life ^1^.

Assembly of PSII is a stepwise process involving multiple accessory factors, many conserved in cyanobacteria and chloroplasts, that promote and regulate biogenesis (Supplementary Fig. 1) ^2^. During the early-stage of PSII assembly in cyanobacteria, the Ycf48 accessory factor binds to the luminal side of the D1 and D2 reaction center complex (RC), preventing premature binding of ions to the Mn_4_CaO_5_ cluster binding site ^3–6^. The PSII RC then binds the CP47 module (CP47_mod_) to form the RC47 complex, followed by attachment of the CP43 module (CP43_mod_) to create an inactive non-oxygen-evolving monomeric PSII complex ^7,8^. Subsequently, fully functional PSII dimers are formed by assembly of the Mn_4_CaO_5_ cluster in a light-driven process known as photoactivation ^9^ and by attachment of extrinsic subunits to the luminal side of PSII to protect and stabilize the Mn_4_CaO_5_ cluster ^10^. At what stage PSII complexes dimerize is still unclear.

Light not only drives photosynthesis and photoactivation but also damages the photosynthetic complexes leading to chronic photoinhibition ^11^. Acceptor- and donor-side mechanisms of visible-light photodamage are still under debate ^11,12^. The acceptor-side mechanism suggests that photodamage results from an over reduced acceptor-side of PSII and blockage of forward electron transfer, which leads to the production of reactive oxygen species, especially singlet oxygen ^11,13,14^. In contrast, donor-side mechanisms propose that excitation of Mn^3+^/Mn^4+^ ions by visible light or the production of reactive oxygen species (ROS) by the Mn_4_CaO_5_ cluster leads to photodamage ^12,15–17^.

Recently, cryo-EM has provided structural insights into various cyanobacterial PSII assembly intermediates, including the PSII RC ^6^, RC47 monomers ^18,19^ and a Psb27/PSII dimer ^20^. These structures were obtained from mutants with deletions of specific PSII subunits that block assembly of PSII to enhance the accumulation of low-abundance PSII intermediates ^6,19,20^. Additionally, structures of PSII lacking the Mn_4_CaO_5_ cluster were obtained by treating oxygen-evolving PSII with NH_2_OH, EDTA and NaCl ^21,22^. The PSII structure missing both the Mn_4_CaO_5_ cluster and extrinsic subunits is termed apo-PSII ^22^.

Previous work by Nowaczyk and colleagues revealed that purification of His-tagged PSII from WT *Thermosynechococcus vestitus* BP-1 *(T. vestitus)* yielded four fractions when separated by anion-exchange chromatography ^23^: a Psb27/PSII monomer, an active PSII monomer, an active PSII dimer and, the fourth, a less active oxygen-evolving PSII dimer fraction, here termed PSII_peak4_, which has not been fully characterized.

Here we have used cryo-EM to determine the structures of the dimeric complexes present in PSII_peak4_ at a resolution of 2.2 Å. This pool of PSII dimers consists of active PSII dimers with extrinsic subunits (PsbO, PsbU and PsbV) bound to both monomers, semi-active PSII dimers with extrinsic subunits attached to only one monomer and inactive dimers that lack extrinsic subunits. Although lacking extrinsic subunits, the inactive PSII monomers in the semi-active and inactive dimers contain all the transmembrane subunits including PsbJ with partial occupancy and all the key cofactors apart from the Mn_4_CaO_5_ cluster and one of the two chloride anions, termed Cl-2. Compared with fully functional PSII, the inactive PSII monomers also show structural differences in D1, D2, CP43 and CP47 around the donor-side close to the binding site of the Mn_4_CaO_5_ cluster. These changes create a channel that allows Mn^2+^ and Ca^2+^ ions access to and from the binding site of the Mn_4_CaO_5_ cluster. In addition, the structure of the D1-His332 residue in the inactive monomer, which is a ligand to the Mn cluster, suggests oxidation of the imidazole side chain. This oxidation could potentially reflect an early stage of photodamage to PSII on the donor-side of PSII.

## RESULTS

### Purification and properties of PSII_peak4_

PSII_peak4_ was purified from *T. vestitus* using His-tagged CP47, as described in ^24^. Consistent with the findings of Nowaczyk and colleagues, PSII_peak4_ is a less active PSII dimer fraction ^23^ (Supplementary Fig. 2; Supplementary Fig. 3). SDS-PAGE with Coomassie brilliant blue staining revealed reduced amounts of the three extrinsic subunits in PSII_peak4_ compared to fully functional PSII (Supplementary Fig. 2; Supplementary Fig. 3), in line with the lower oxygen evolution activity of PSII_peak4_. It is unlikely that the extrinsic subunits were removed by salt during anion exchange, as the elution buffer contained 5-200 mM MgSO_4_ ^24^, which is insufficient to detach extrinsic subunits ^25^. Negative staining electron microscopy confirmed that PSII_peak4_ contained dimeric PSII complexes similar in size to fully functional PSII (Supplementary Fig. 4).

### Overall structures of PSII_peak4_ intermediates

The composition and structures of the PSII complexes in PSII_peak4_ were determined by cryo-EM single particle analysis. The dataset was collected with the grid tilted 30 degrees to mitigate the effects of preferential orientation of the sample. Extensive 2D and 3D classification resulted in 1.06 million good particles which were separated into three classes: active PSII dimer (34.5% of total particles), inactive PSII dimer (39.6% of total particles) and semi-active PSII dimer (25.8% of total particles) (Fig. 1, Supplementary Fig. 5, Supplementary Fig. 6). The presence of inactive complexes explains the reduced oxygen-evolution activity in PSII_peak4_ (Supplementary Fig. 3c). The distinction between these classes is based on the presence or absence of the extrinsic subunits (PsbO, PsbU and PsbV) and the Mn_4_CaO_5_ cluster on the luminal side of PSII. Active dimers contain both sets of extrinsic subunits, inactive dimers lack both sets of extrinsic subunits, and semi-active dimers have one active monomer and one inactive monomer lacking the extrinsic subunits. The overall resolution for all three classes is 2.2 Å. The data processing pipeline is outlined in Supplementary Fig. 6 and the refinement statistics are shown in Supplementary Table 1. Examples of the density fit are shown in Supplementary Figures 7-10 and statistics of modelling are shown in Supplementary Tables 2-4.

**Figure 1.**
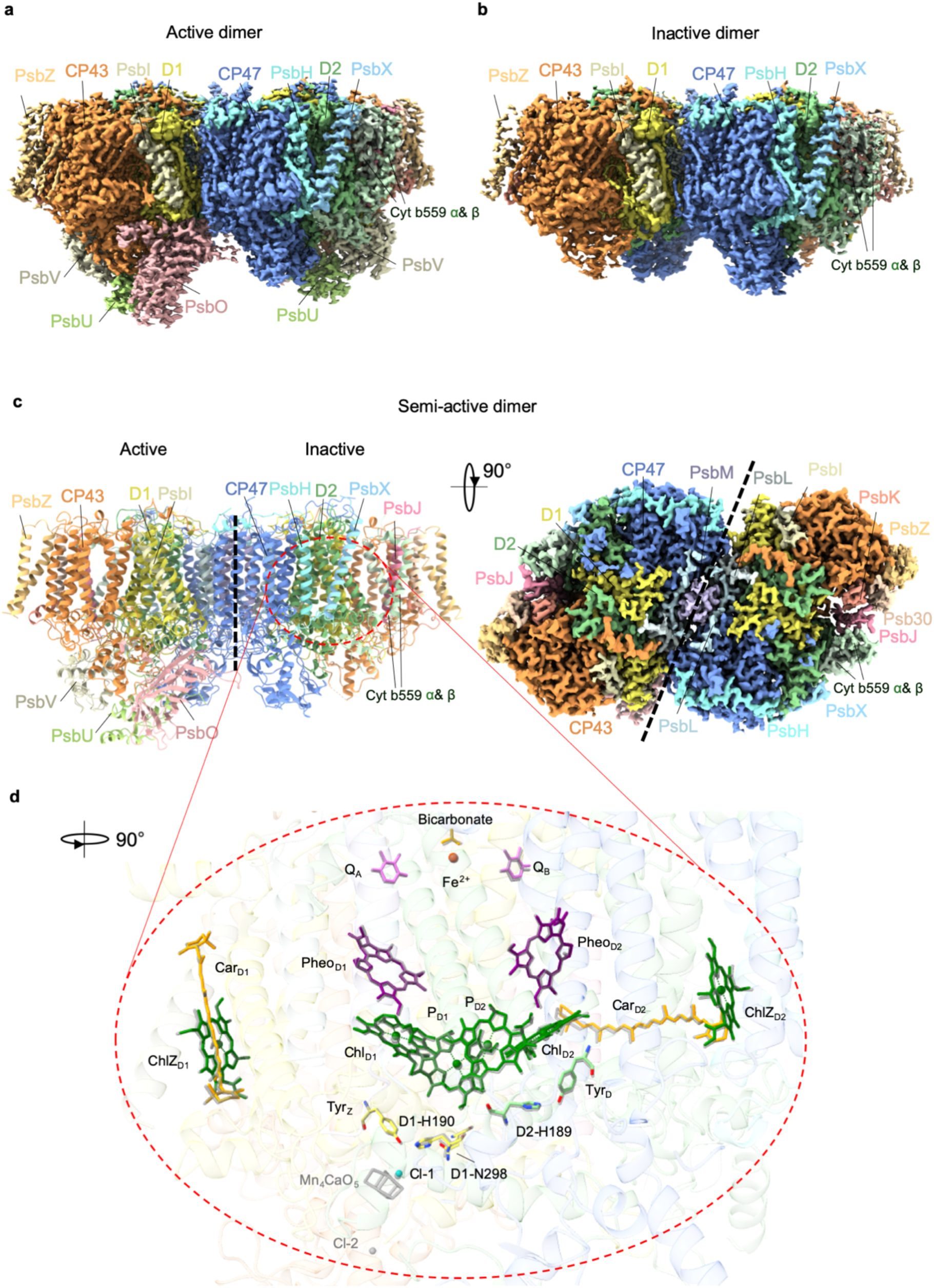
The cryo-EM maps and models of PSII_peak4_ complexes. **a.** Active dimer map. **b**. Inactive dimer map. **c.** The structure and map of the semi-active dimer. **d.** Comparison of cofactors in the inactive PSII and fully functional PSII. The cofactors of fully functional PSII are shown in grey and the inactive PSII cofactors are shown in color.

The active monomers in the active and semi-active dimers contain the D1, D2, CP43 and CP47 subunits, 12 low-molecular-mass transmembrane subunits PsbH, PsbX, Cyt b559 α&β, PsbI, PsbK, Psb30, PsbZ, PsbL, PsbT, PsbM and PsbJ, and three extrinsic subunits, PsbO, PsbU and PsbV (Fig. 1 a,b,c). The active monomer shows high similarities to the high-resolution crystal structure of *Thermosynechococcus vulcanus* (*T. vulcanus*) PSII ^26^ with a root mean square deviation (rmsd) of 0.28 Å. The inactive monomers (lacking the Mn_4_CaO_5_ cluster and extrinsic subunits) contain all 16 transmembrane subunits and are highly similar to the transmembrane portion of fully functional PSII, although the single helix PsbJ subunit, involved in the late stage of PSII assembly ^8^, had a lower EM density due to either reduced occupancy or higher flexibility (Fig. 1c). On the luminal side, the inactive PSII complex lacks not only the extrinsic subunits and the Mn_4_CaO_5_ cluster, but also the Cl-2 ion and density for the D1 C-terminal tail from D1-N335 to A344, as well as the luminal region of CP43 from G313 to L325 (Fig. 1d) (Supplementary Table 2, 4).

The inactive monomer in the semi-active dimer closely resembles those present in the inactive dimer with a rmsd less than 0.32 Å. Similarly, the active monomer in the semi-active dimer shows a rmsd less than 0.21 Å compared to those in the active dimer. Therefore, subsequent structural descriptions and comparisons are based on the inactive and active monomers from the semi-active dimer PSII, unless stated otherwise.

### Cofactors in the inactive monomer

In the active PSII complex, the arrangement of key cofactors is highly similar to that of fully functional PSII determined previously (Supplementary Fig. 11). In the inactive PSII, apart from the Mn_4_CaO_5_ cluster and Cl-2, other key cofactors and residues from D1/D2 heterodimer are all present at their expected positions, including P_D1_, P_D2_, Chl_D1_, Chl_D2_, Pheo_D1_, Pheo_D2_, Q_A_, Q_B_, the non-heme iron, bicarbonate, D1-β-carotene, D2-β-carotene, ChlZ_D1_, ChlZ_D2_, Cl-1 (Fig. 1) and the chlorophylls and β-carotenes in CP43 and CP47 (Supplementary Fig. 12). It is known that the Mn_4_CaO_5_ cluster is not stable without the extrinsic subunits ^27^ and that Cl-2, proposed to be involved in a proton-exit pathway, binds to D1-Asn338, D1-Phe339 and CP43-Glu354 (Glu342 in *T. vestitus*) close to the interface of D1 and the extrinsic subunits ^26,28,29^. Therefore, the absence of the Cl-2 is likely associated with the lack of a folded D1 C-terminal tail (Fig. 2c).

**Figure 2.**
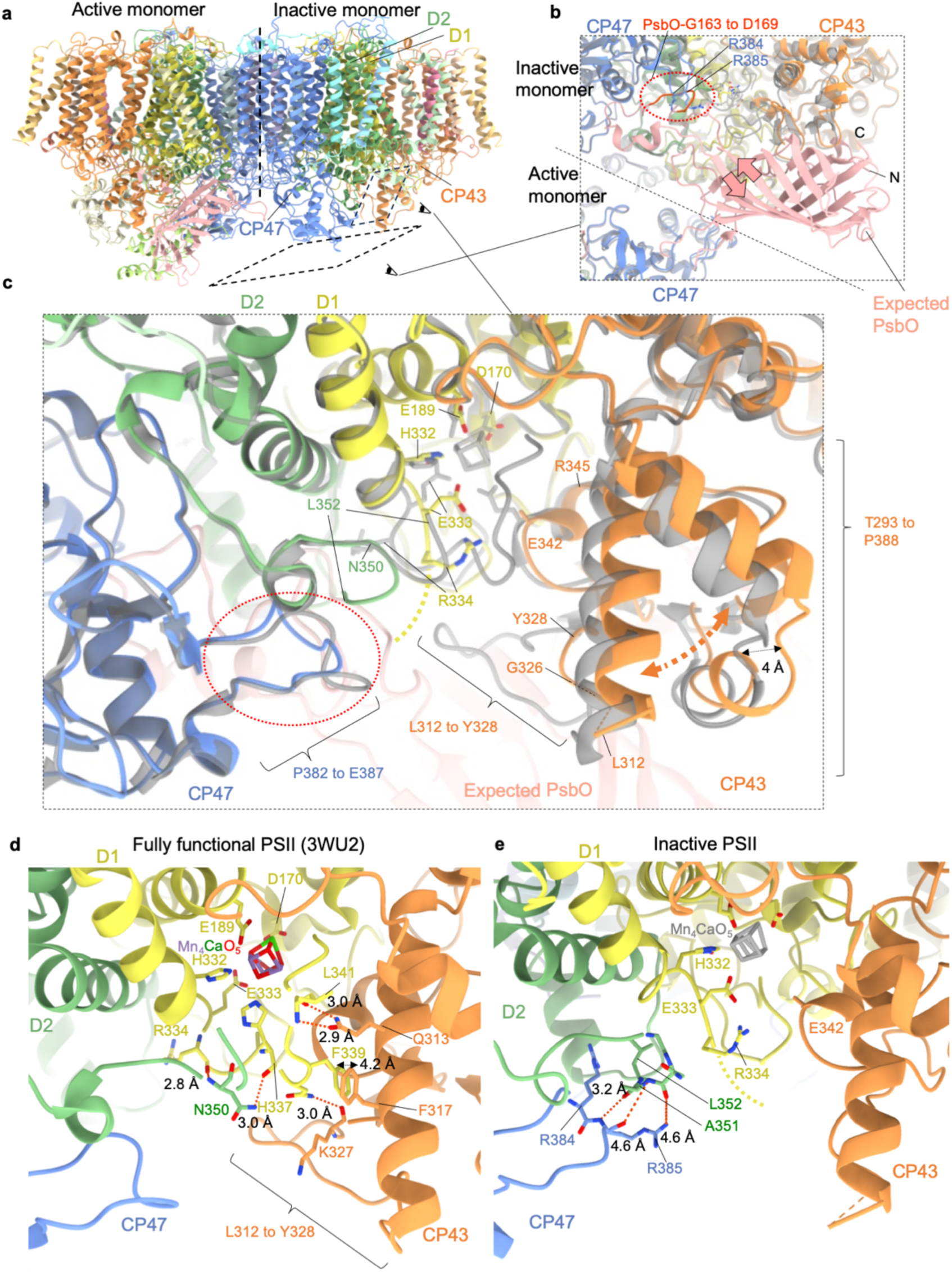
Structural differences between the fully functional PSII and inactive monomer at luminal region. **a.** Structure of the semi-active PSII with some subunits labelled for inactive monomer. **b.** Large external loop of PsbO from fully functional PSII clashes with CP47 in the inactive monomer. The red dashed circle in (**b, c**) indicates the displacement of CP47 and the clashes with the expected binding site of PsbO. The region of PsbO which clashes with CP47 is highlighted in red. The PsbO loop from N- to C-terminus is indicated by arrows. **c.** Overlay of the fully functional (gray) and inactive PSII monomer (colored). **d, e.** Detailed interactions in the CP43-L312 to Y328 region of the fully functional (**d**) and inactive PSII monomer (**e**). The active PSII structure was taken from *T. vulcanus* crystal PSII structure (PDB ID: 3WU2). The CP43 of fully functional PSII was renumbered to match the residue number in *T. vestitus*. The residues shown in the figure are conserved in *T. vulcanus* and *T. vestitus*.

Redox-active tyrosines Y_Z_ (D1-Tyr161) and Y_D_ (D2-Tyr160), along with the related H-bonding histidine residues D1-His190 and D2-His189, are located at the expected positions (Fig. 1d). The distance between Y_z_ and His190 is crucial for the rapid reduction of P_D1_ ^+^ ^30^. In the inactive monomer, this distance is 2.5 Å, which is close to the value of 2.4 Å in oxygen-evolving PSII ^26^. Additionally, water molecules, which contribute to shortening the distance between Tyr_Z_ and His190 ^30^, are observed in similar positions in the inactive and active monomers (Supplementary Fig. 13). D1-Asn298, which is H-bonded to His190 and is necessary for efficient oxidation of Tyr_Z_ ^30^, is also located at the expected position (Fig. 1d, Supplementary Fig. 13).

In general, our inactive PSII complexes display well-resolved cofactor density on the acceptor side, including Q_B_ (Supplementary Fig. 10), and closely resemble the structure of fully functional PSII in this region (Supplementary Fig. 14), while lacking cofactors on the donor side (Fig. 1d).

### The extrinsic proteins are not required for accumulation of dimeric PSII

Previous studies of PSII from the mesophilic cyanobacterium *Synechocystis* sp. PCC 6803 have suggested that PsbO stabilizes the dimeric state of PSII ^22,31^. The lack of PsbO in the inactive PSII dimer and Psb27/PSII dimer ^20^ would suggest that PsbO is not obligatory for dimerization and that attachment of extrinsic subunits to the two monomers in a dimer may occur independently.

Comparison of the structures allowed us to investigate the role of the extrinsic proteins in modifying the structures of the intrinsic subunits. Interestingly, attachment of PsbO in the active monomer results in a shift of a minor loop from CP47-G82 to CP47-P88 in the CP47 subunit in the inactive monomer (Supplementary Fig. 15). Apart from this difference in the CP47 loop, no other structural changes were observed at the interface that could be attributed to the attachment of the extrinsic subunits.

We found here that a ß-sheet in CP47 (CP47-H343 to R347), previously reported to be disordered upon removal of extrinsic proteins ^22^, remains ordered and identical to that of the fully functional PSII in the active PSII monomer from semi-active PSII dimer, despite the absence of the adjacent PsbO (Supplementary Fig. 16). The disordered ß-sheet in this region is therefore likely caused by loss of low-molecular-mass subunits, PsbJ, PsbY and PsbZ in apo-PSII ^22^.

### Presence of an open channel to the Mn-binding site in inactive complexes

Compared with the fully functional PSII complex, the inactive PSII complex exhibits not only the absence of extrinsic subunits but also structural differences on the luminal side. In fully functional PSII, the Mn_4_CaO_5_ cluster is covered by extrinsic subunits as well as the C-terminal tail of D1 and D2 and a luminal loop of CP43 from L312 to Y328 (Fig. 2 c, d). In contrast, structural changes in the inactive PSII complex, including alterations in the luminal region of CP43, the C-terminal tail of D1 and D2, and a minor loop of CP47, result in the opening of a channel reaching from the solvent to the site of binding of the Mn_4_CaO_5_ cluster (Fig. 2 c, e). This negatively charged channel has a radius of 2.1 Å at its narrowest point, which is larger than the atomic radii of Mn^2+^ (1.6 Å) and Ca^2+^ (1.9 Å) (Supplementary Fig. 17).

### CP43 displacement and flexibility prevents binding of extrinsic subunits

In the inactive PSII complex, the luminal region of CP43 from T293 to P388 shifts away from the Mn_4_CaO_5_ cluster with the most prominent displacement (∼4 Å) observed between CP43-D364 and CP43-A380. However, CP43-E342 (E354 in *T. vulcanus*), which acts as a ligand to the Mn_4_CaO_5_ cluster, is present at the expected position (Fig. 2c). The density of the side chain of CP43-E342 cannot be resolved clearly, suggesting different rotamers.

A previous atomic force microscopy (AFM) study of *T. vulcanus* suggested that assembly of the Mn_4_CaO_5_ cluster induces changes in the positions of CP43-E354 and CP43-R357, subsequently triggering conformational changes in the luminal region of CP43 ^10^. While our data supports a shift in this region of CP43, it is of a significantly lower magnitude, and the corresponding residues implicated in triggering conformational changes (E342 and R345 in *T. vestitus*) remain at the positions observed in fully functional PSII (Fig. 2c).

Similar shifts in CP43 from T293 to P388 have been reported in other PSII intermediates, including apo-PSII, the Psb27/PSII dimer and Psb28/Psb27/PSII ^18,20,22^. The shift in this area does not cause steric clashes with the expected positions of PsbO, PsbU, or PsbV (Supplementary Fig. 18). Instead, the displacement of the CP43 luminal domain shifts it away from the Mn_4_CaO_5_ cluster, resulting in the disengagement of CP43 from the binding interface for the extrinsic subunits.

Additionally, the luminal loop CP43-G313 to L325 in the inactive monomer exhibits poor density and cannot be modelled, suggesting high flexibility in this region (Fig. 2c, Supplementary Fig. 19). Furthermore, CP43-G326 to Y328 region shows an alternative orientation to that observed in fully functional PSII so that CP43-K327 points away from the D1 C-terminal tail (Fig. 2c). Other PSII intermediates lacking the extrinsic subunits also lack density at this area, similar to our inactive PSII monomer (Supplementary Fig. 20) ^18–20,22^.

In agreement with Gisriel and colleagues ^22^, the flexibility of CP43 is likely due to the absence of extrinsic subunits. However, we notice two key differences between CP43 in the inactive PSII complex and Gisriel’s apo-PSII structure ^22^. First, CP43-E354, which shifts 3 Å away from its expected Mn coordinating position in apo-PSII, remains unchanged in our structure. Second, while the region from CP43-G389 to F404 is difficult to model in apo-PSII ^22^, most of this region exhibits clear density in our structure. Possibly the loss of low-molecular-mass subunits, PsbJ, PsbY and PsbZ, in apo-PSII might increase the flexibility of the soluble domain of CP43.

### Flexible D1 C-terminal tail

Although the Mn_4_CaO_5_ cluster-coordinating residues D1-D170 and D1-E189 remain at the expected position in the inactive PSII complexes, D1-E333 and D1-R334 show a different conformation to that in fully active PSII (Fig. 2c). D1-E333 undergoes a ∼ 4 Å shift, with its side chain orientated toward the Mn_4_CaO_5_ cluster, in an opposite orientation to that modelled in apo-PSII ^22^ (Supplementary Fig. 21). Density corresponding to the D1 C-terminal tail from N335 to A344 could not be observed in the inactive monomer likely due to the high flexibility in this region (Fig. 2) as mass spectrometry data confirmed that the D1 C-terminus was not truncated (Fig. 3b).

**Figure 3.**
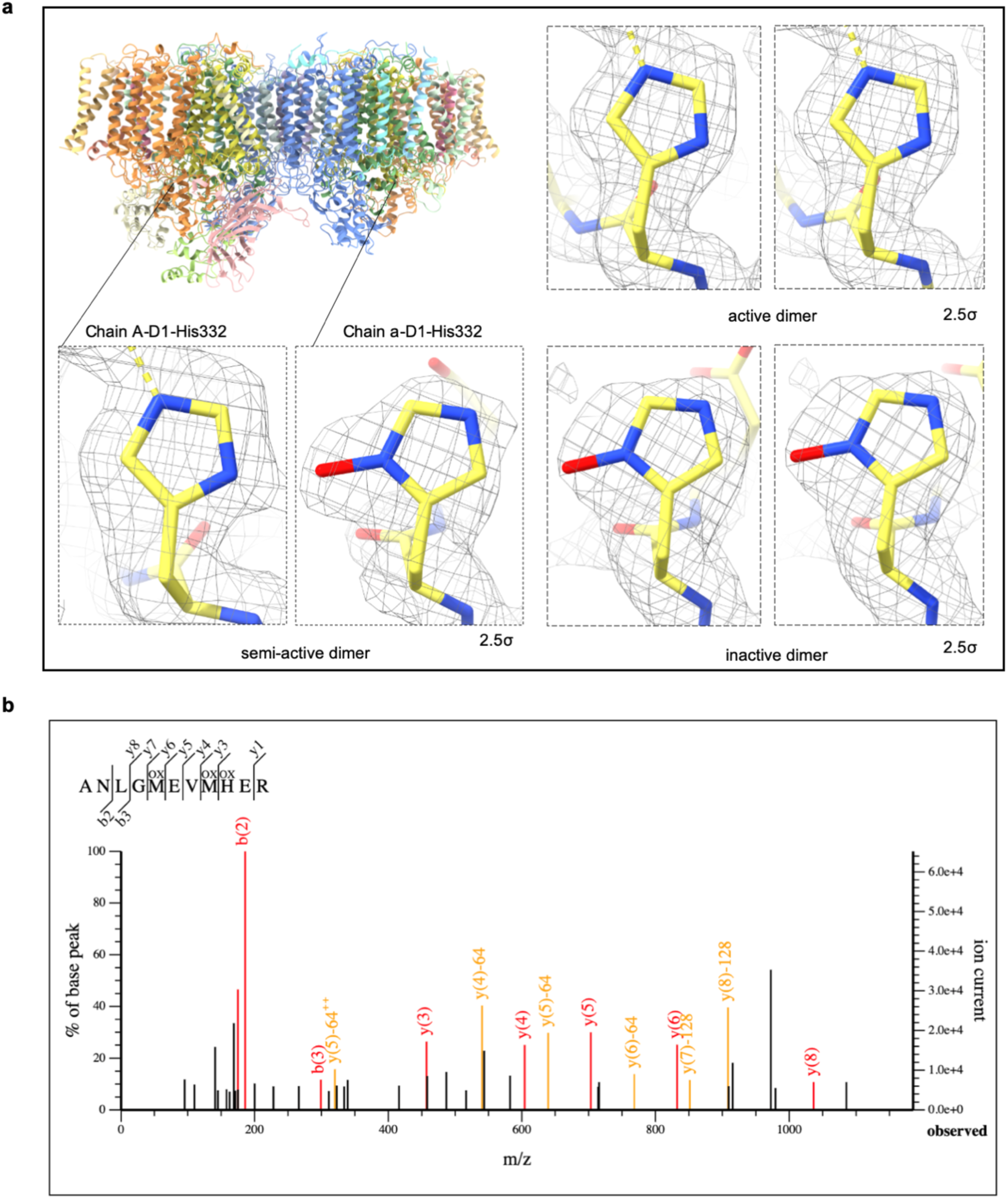
Oxidation of D1-H332 in the inactive PSII monomer. **a,** Density of D1-His332 at the inactive, semi-active and active dimer. Densities are shown at the contour level of 2σ. **b**, Mass spectrometry of oxidation of the C-terminal tail of D1. The whole peptide has a mass of 48 Da greater than non-oxidized peptide, so it has three oxidations in total. y(4) and y(4)-64 confirm M331 is single oxidized, y(8) and y(8)-128 confirm M228 is also single oxidized. Thus H332 is oxidized. Software didn’t compare to the non-oxidized peptide so there is no y(3)-16 peptide on the figure.

Previous structures of PSII intermediates have also identified D1-D170 and D1-E189 at the Mn coordinating positions ^6,18–20,22^. Notably, D1-170 and D1-E189 are already positioned to bind the Mn cluster in the early reaction center assembly intermediate (RCII) ^6^ with D1-D170, coordinating Mn_4_, previously identified as a high-affinity Mn^2+^ binding site ^32^. Additionally, a Psb27/Psb28/PSII assembly complex ^18^ contained an unknown metal ion density close to D1-Asp170. These observations further support the hypothesis that the primary high-affinity Mn^2+^ binding site is close to Mn_4_ and Ca^2+^ ^22,33^.

Unexpectedly, the side chain of D1-His332 in the inactive monomer was found to have an extra density in addition to the imidazole ring (Fig. 3a). Mass spectrometry data confirmed that D1-H332 was single oxidized (Fig. 3b), so we propose that this density arises from the oxidation of the imidazole ring to form hydroxylated histidine, similar to that previously reported for D2-His336 ^34^. To fit this density, D1-H332 needs to adopt a flipped rotamer from its Mn-coordinating conformation. The possibility that the additional density reflects the presence of different rotamers of D1-H332 is unlikely, as different rotamers reduce density occupancy and the imidazole ring cannot be fully aligned with the extra density (Supplementary Fig. 22). The 2.2 Å resolution of the model is insufficient to resolve protonation as an explanation for the extra density, and the density is too close to the imidazole ring to be due to a ligated metal ion^35^. In apo-PSII, D1-H332 was found to be close to Cl-1 ^22^, but in our structure, this histidine is approximately 4 Å away from Cl-1 and so unable to coordinate it.

One possibility to explain possible oxidation of D1-H332 is that D1-H332 undergoes photodamage due to oxidation by ROS generated during water oxidation and that the inactive complexes represent damaged complexes rather than assembly complexes. D1-His332 is close to the site of water oxidation as it ligates Mn_1_ of the cluster which is thought to bind the O_x_ water involved in the formation of O_2_ through an oxo-oxyl coupling mechanism ^29^, possibly involving a peroxide intermediate ^36^. Although 2-oxo-histidine appears to be more commonly observed in biology under ROS-mediated conditions ^37,38^ than hydroxylated histidine (Supplementary Fig. 23), the specific oxidation product likely depends on the local protein environment and structure ^39^. Early photoinduced oxidation of D1-H332 has also been observed in experiments using isolated PSII complexes ^40,41^.

In addition, single methionine oxidation was found by mass spectrometry in the D1 C-terminal tail (M328 and M331). However, apart from a slight extra density at D1-M331 in active dimers, no clear additional density was found on the side chain of D1-M328 and M331 in either active or inactive PSII (Supplementary Fig. 24). Artefactual single oxidation of Met has previously been observed during the preparation of mass spectrometry samples ^42^, whereas, to our knowledge, single oxidation of histidine residues has not been observed under similar conditions.

### D1 C-terminus unfolding allows flipping of D2 C-terminus and displacement of CP47

In oxygen-evolving PSII, the D1 C-terminal tail not only provides ligands (H332, E333, D342, A344) to the Mn_4_CaO_5_ cluster, but also interacts with the C-terminal tail of D2, the CP43 L312-Y328 loop, and a helix from T293 to G313 of CP43 (Fig. 2d). The backbone oxygens of D1-R334 and D1-H337 H-bond to the backbone nitrogen and side chain of D2-N350, respectively. The side chain of D1-N338 H-bonds to the backbone of CP43-K327. In addition, D1-F339 exhibits a hydrophobic interaction with CP43-F317, and CP43-Q313 also H-bonds to D1-L341.

In the absence of a structured D1 C-terminal tail, the C-terminal tail of D2 flips to the opposite orientation, interacting with CP47-R384 within a small loop of CP47 (Fig. 2c, e). This interaction induces a minor shift in the loop of CP47 and results in clashes with PsbO-G163 to D169 in the large external loop of PsbO (Fig. 2b), a crucial region for the attachment of PsbO ^43^. Interestingly, mutation of CP47-R384 and R385 decreases the binding of extrinsic subunits ^44^, suggesting that this interaction between this C-terminal tail of D2 and CP47 is important for assembly and possibly disassembly of PSII.

## DISCUSSION

We have determined the structures of inactive, semi-active and active dimeric PSII complexes at a resolution of 2.2 Å from *T. vestitus*. The semi-active and inactive PSII may represent either assembly intermediates or photodamaged disassembly complexes. The observed oxidation of the D1-His332 residue, a ligand of the Mn cluster, suggests that the PSII_peak4_ population is more likely derived from photodamaged PSII. The naturally occurring PSII_Peak4_ complex represents about 5% of total PSII (Supplementary Fig. 2), suggesting it is a stable intermediate in wild-type *T. vestitus*. This implies it could be a pool of backup PSII dimers prepared for rapid replacement of damaged PSII or that PSII_peak4_ are damaged PSII complexes waiting for disassembly and repair. From a physiological perspective, prompt detachment of the extrinsic proteins triggered by photodamage to D1-H332 would allow access of reductant to the Mn cluster to promote disassembly ^45^ thereby preventing the formation of further reactive oxygen species.

A semi-active PSII dimer has also been observed after affinity purification of twin-strep tagged PsbO followed by anion-exchange chromatography. This study suggested that the partial oxygen-evolving activity may result from incomplete occupancy of the Mn₄CaO₅ cluster ^46^.

A recent structure of a Psb32/PsbV/Psb27/PSII assembly intermediate has revealed the binding of the Psb27 and Psb32 assembly factors, along with the PsbV extrinsic subunit, to the luminal side of PSII ^47^. In this complex, the transmembrane helix of Psb32 occupies the position of PsbY and assists in binding of PsbV, suggesting that extrinsic subunits are not incorporated spontaneously, but rather in a sequential and regulated manner with assistance of Psb32.

In contrast, our inactive PSII structures show clear density for PsbY but lack all extrinsic subunits, supporting the idea that the Peak4 complexes may derive from photodamaged PSII. However, the possibility that they represent late-stage assembly intermediates cannot be excluded, if multiple pathways exist for PSII assembly. This is supported by the observation that *psb32* deletion mutants of cyanobacteria and plants exhibit phenotypic deficiencies only under stress conditions ^48,49^, implying that Psb32 is not absolutely essential for PSII assembly and alternative pathways may exist.

Overall, on the basis of the structural differences between the inactive and active PSII complexes, we propose a mechanism (Fig. 4) for the assembly/disassembly of oxygen-evolving PSII. Furthermore, because of the similarities between PSII assembly and disassembly intermediates, we suggest that these processes follow similar trajectories but proceed in opposite directions.

**Figure 4.**
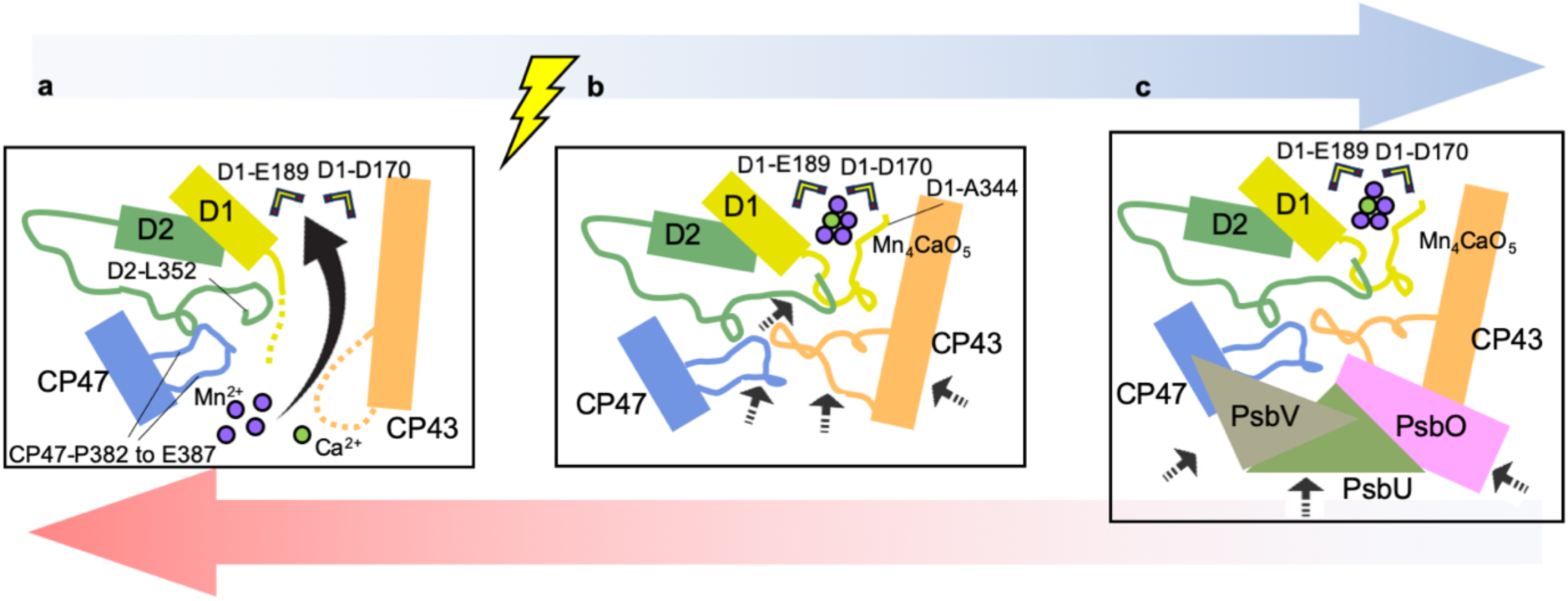
The mechanism of the final stages of PSII assembly. Cartoon panels from **a** to **c** represent assembly pathway (blue gradient arrow), while **c** to **a** sequence represents disassembly pathway (red gradient arrow). Mn^2+^ and Ca^2+^ ions are initially delivered to the binding site of the Mn_4_CaO_5_ cluster. Photoactivation then drives the assembly of the Mn_4_CaO_5_ cluster and the folding back of the D1 C-terminal tail. The folding of D1 induces the flipping of D2 C-terminal tail, structural rearrangements in the CP47 loop, and movement of the CP43 loop toward the Mn_4_CaO_5_ cluster. Finally, the extrinsic subunits attach to the luminal side of the PSII, completing the formation of fully functional PSII. The attachment of extrinsic subunits may not occur spontaneously but rather in a sequential manner. PsbV may bind to the luminal side early in the process, while PsbU is likely incorporated at a later stage.

We propose that photoactivation which drives the assembly of the Mn cluster is coupled to the remodeling of the C-terminal tail of D1 within the complex. The folded D1 triggers the movement of the luminal domain of CP43 and releases the CP47 loop from D2 C-terminal tail to allow the attachment of the PsbO extrinsic subunit (Fig. 4). The current two-quantum photoactivation model, based largely on biochemical data, fits well with our mechanism ^9^. After the first flash oxidizing a Mn^2+^ ion bound to the high-affinity Mn-binding site, PSII undergoes a slow ‘dark rearrangement’ that possibly corresponds to the movement of D1, D2, CP43 and CP47 in the luminal region. A second photon then oxidizes a second Mn^2+^ ion to generate the first stable intermediate in assembly of the Mn cluster.

Our structure of the inactive PSII complex revealed the presence of the putative channel for the delivery of metal ions for Mn cluster assembly (Supplementary Fig. 17). Gisriel et al suggested that D1-E333 in the C-terminal tail could deliver Mn^2+^ to the Mn cluster binding site ^22^. However, this remains an open question as our structure shows an opposite orientation of D1-E333 to that of apo-PSII (Supplementary Fig. 21). It is feasible that one of the water channels, the O1 channel, which has a minimum diameter exceeding 1.7 Å, slightly larger than the atomic radius of Mn^2+^ (1.6 Å) ^50^, may also serve as a Mn delivery channel, assuming Mn is dehydrated in the channel. However, mutations of O1 channel bottleneck residues do not appear to affect the assembly or structure of the Mn₄CaO₅ cluster ^51^.

Inactive PSII complexes show Q_B_ density (Supplementary Fig. 10), while lacking cofactors on the donor side (Fig. 1d). This observation is similar to the recently reported Psb32/Psb27/PsbV/PSII assembly intermediate ^47^. In addition, the acceptor-side of inactive PSII resembles that of the structure of fully functional PSII (Supplementary Fig. 14). These observations suggest that the acceptor side is structured prior to the donor side of PSII during assembly, and that disassembly likely initiates on the donor side, triggering subsequent breakdown of the entire complex. This may prevent blockage of the electron flow and subsequent photodamage to PSII intermediates ^11^.

The flexible C-terminal tail of D1 is a feature of all PSII intermediates ^19,20,22^, apart from the Psb27/Psb28/PSII complex which shows an unknown metal bound to the luminal side of PSII ^18^ (Supplementary Fig. 21). For all structures of PSII intermediates, the re-orientated D2 C-terminal tail is present (Supplementary Fig. 21), but density for the loop from CP43-G313 to L325 is always missing, even though in some cases this region was still built in the model (Supplementary Fig. 20) ^18–20^.

The structure of the soluble domain of CP43 in the inactive PSII monomer is highly similar to that of the *T. vulcanus* Psb27/PSII dimer ^20^ and *T. vestitus* Psb28/Psb27/PSII monomer ^18^ (Supplementary Fig. 25). When our inactive monomer is superimposed on the two Psb27-containing structures ^18,20^, only minor clashes are observed between Psb27 and our CP43 (Supplementary Fig. 25). Additionally, several CP43 residues previously reported to conflict with and disrupt interactions with PsbO were found in similar positions in our inactive PSII ^20^ (Supplementary Fig. 26). These observations suggest that the existence of Psb27 is unlikely to induce structural changes in CP43. Instead, Psb27 likely acts as a ‘door wedge’, blocking the necessary movement of CP43 required for Mn_4_CaO_5_ cluster assembly. If the soluble domain of CP43 from the Psb27/PSII structure is moved to its position in the fully functional PSII, Psb27 sterically clashes with PsbO (Supplementary Fig. 27) ^20^, indicating that with Psb27 bound, the soluble domain of CP43 cannot shift toward the Mn_4_CaO_5_ cluster and properly accommodate PsbO. As a result, the proper ‘dark rearrangement’ during photoactivation may be impeded.

## METHODS

### Purification and characterization of Peak 4 PSII

The His tagged CP47 *T. vestitus* cells were grown under ∼60 μmol photons·m^−2^·s^−1^ light intensity, and thylakoid membranes were solubilized using 1% DDM for 10 min in 4 ℃ at chlorophyll concentration of 1 mg/ml. The solubilized PSII was purified by nickel affinity chromatography followed by anion exchange chromatography according to Nowaczyk et al ^23^ with the following modifications: PSII complexes eluted from nickel affinity column were loaded directly on the Bio-Rad UNO Q-12 column using a AKTA purifier 10 system; PSII were then eluted with 5-200 mM MgSO_4_ gradient ^24^. The purification methods were derived from ^23^ and described in ^24^.

### Measurement of oxygen evolution activity

Oxygen evolution activity was assessed using an Oxyl1-Oxylab device (Hansatech Instruments, UK). Cells were adjusted to a chlorophyll a (Chl a) concentration of 1 μg/ml. The activity was measured with 1 mM potassium ferricyanide (KFeCN) and 0.1 mM 2,6-dichlorobenzoquinone (DCBQ) under a light intensity of 2000 μmol photons m⁻² s⁻¹. For PSII protein samples, 1 μg of Chl a was mixed with 1 ml of a buffer solution containing 50 mM MES-NaOH (pH 6.5), 500 mM sucrose, 30 mM CaCl₂, 10 mM MgCl₂, 1 mM KFeCN, and 0.1 mM DCBQ. The oxygen evolution activity was then measured under the same light conditions of 2000 μmol photons m⁻² s⁻¹.

### Mass spectrometry

D1-His332 (P0A444; PSBA1) is contained within the D1 peptide residues 324 – 334, ANLGMEVMHER, and consequently it is necessary to distinguish histidine oxidation from methionine oxidation events. PSII was digested with trypsin and peptides were analyzed by nano-liquid chromatography with data-dependent high-resolution tandem mass spectrometry (nLC-MS/MS) that provides mass information within the peptide to residue-specific resolution. Data was matched to the *T. vestitus* proteome using Mascot software with variable modification of methionine residues to either sulfoxide (+15.9949 Da) or sulfone (+31.9898 Da) and/or histidine residues to single oxidized histidine species (+15.9949 Da). Unmodified and peptides with varying levels of oxidation were identified in the chromatogram. Histidine oxidation was identified in peptides with single or double Met oxidation (Met328 & Met 331) with y-ion fragments providing residue-specific resolution.

### Grid preparation

The sample, concentrated 0.3 mg/ml in 20 mM MES pH 6.5, 2.5 mM MgCl_2_, 2.5 mM CaCl_2_, 7 mM MgSO_4_, 0.03% DDM, 0.05% CHAPS, was frozen on Quantifoil 0.6/1, 300 mesh copper grids manually coated with a continuous layer of carbon approximately 1 nm thick ^52^. The grids were glow-discharged for 5 seconds at 25 mA prior to freezing. The sample was prepared using a Vitrobot Mark IV, set to 4°C, 100% humidity, 2” blot time and 25 blot force. 3 µl of sample were applied to each grid. The sample was manipulated under green light until frozen.

### Data collection

The micrographs were collected on a 300 kV Titian Krios G3i, equipped with Gatan K3 BioQuantum direct detector. Preliminary data collection indicated that, although showing good particle spread, the sample adopts preferential orientation on continuous carbon. Therefore, the high-resolution dataset was collected with grid tilted 30 degrees, to mitigate the effects of preferential orientation. As shown in Supplementary Fig. 6, about 9173 movies were collected in 72 hours, using the software Serial-EM for 48 hours and EPU for 24 hours. The grid was aligned so that rows of holes would sit along the tilt axis: this allowed to collect 9 holes with beam-image shift while keeping the beam tilt below 10 mrad ^53^. The data collection details are summarized in Supplementary Table 1.

### Data processing

The processing outline is described in Supplementary Fig. 6. After optics group assignment, the raw micrographs were aligned using the own implementation of motion correction from Relion 3.1 ^54^. CTFFIND ^55^ was used for initial CTF estimation, then Relion 3.1 was used for particle picking, 2D and 3D classification, polishing and CTF/3D refinements, except for per-particle defoci, which were estimated using GCTF ^56^ upon particle picking, to immediately assign the correct defoci to each particle. This step was important, since CTFFIND only estimates per-micrograph defocus, not considering the defocus change resulting from the stage tilt. Initial picking was performed with LoG in Relion, resulting in 5.9 Mln particles, then subjected to 2D classification. 3.4 M good particles were selected, pooled, globally refined, then filtered based on LogLikeliContribution (up to 91300), Autopick Figure of Merit (FoM) (max 0.2), CTF FOM (min 0.1), NormCorrection (0.57-0.61) to remove particles featuring strong artefacts in the micelle area. The resulting 2.65 M particles were polished and CTF-refined, then separated by 3D classification without alignment (20 Å lowpass filter of reference map, T64 and 10 classes), to resolve the compositional heterogeneity (presence vs absence of the Psb O-U-V subunits). Pre-alignment of particles via global refinement and subsequent classification without alignment resulted in better separation than 3D classification with alignment. This strategy allowed to isolate two good classes, without artefactual streaks: a class of 636664 particles, featuring a mixture of active (Psb O-U-V present in both PSII monomers) and semi-active (Psb O-U-V present only in one PSII monomer) particles and one class of 420178 inactive (Psb O-U-V absent) particles. The two classes were then further classified without alignment (20 Å lowpass filter of reference map, T64 and 3 classes) to fully separate the particles and this led to the three final classes: active PSII (363811 particles, 2.4 Å), semi-active PSII (272215 particles, 2.2 Å) and inactive PSII (417537 particles, 2.5 Å). Since the active and inactive maps were perfect dimers, the maps were also postprocessed with C2 symmetry, to further improve resolution, yielding 2.2 Å maps. Post-processing and local resolution estimation were performed in Relion 3.1.

### Model fitting

The crystal structure of *T. vestitus* monomeric PSII (PDB ID: 3KZI)^57^ and cryo-EM structure of *T. vestitus* monomeric PSII lacking PsbJ and extrinsic subunits (PDB ID: 7NHO) ^18^ were placed into cryo-EM density of active and inactive PSII monomers within the structures of PSII dimers using phenix.dock_in_map ^58,59^. The model of PsbJ from monomeric PSII ^57^ was used as the initial model for PsbJ in the inactive PSII monomers. Subsequently, the structures of active and inactive monomers were corrected and adjusted in COOT to match cryo-EM density ^60^. Next, the refinement was carried out using phenix.real_space_refine with cofactor restraints generated by phenix.elbow ^58,59^. Active/inactive monomeric PSII complexes were then merged to create three PSII dimer structures, which were further refined with COOT and phenix.real_space_refine. Residues in the structures were carefully inspected in COOT, and iterative rounds of manual refinement were performed using COOT and phenix.real_space_refine until no significant errors were present in COOT and MolProbity (http://molprobity.biochem.duke.edu/) ^61^ validation reports. Density fitting and refinement statistics are shown in Supplementary Figures 7-10 and Supplementary Tables 2-4.

## Data availability

The cryo-EM maps are deposited in the Electron Microscopy Data Bank under accession number EMD-51100 (inactive dimer), EMD-51102 (active dimer) and EMD-51101 (semi-active dimer). The models are deposited in the Protein Data Bank under accession numbers 9G6F (inactive dimer), 9G6H (active dimer) and 9G6G (semi-active dimer).

Source data are provided within this paper.

## ACKNOWLEDGEMENTS

P.J.N. is grateful for the support of the Biotechnology & Biological Sciences Research Council (awards BB/L003260/1 and BB/P00931X/1). L.A.S. acknowledges the support by the Scientific Service Units (SSU) of IST Austria: the Electron Microscopy Facility (EMF), the Life Science Facility (LSF) and the IST high-performance computing cluster.

## CONTRIBUTIONS

P.J.N., L.A.S., conceived the research; Z.Z., I.V., K.M. and J.P.W. performed the experiments; I.V., Z.Z. and J.P.W. analyzed the data; all authors contributed to writing of the manuscript.

## Abbreviations

PSII: Photosystem II
OEC: Oxygen-evolving complex
RC: D1 and D2 reaction center complex
CP47mod: CP47 module containing PSII subunits CP47, PsbH, PsbL, PsbM, PsbT, PsbX and accessory factors
CP43mod: CP43 module containing PSII subunits CP43, PsbK, Psb30, PsbZ and accessory factors
ROS: Reactive oxygen species
T. vestitus: Thermosynechococcus vestitus BP-1
T. vulcanus: Thermosynechococcus vulcanus
rmsd: root mean square deviation
RCII: Reaction Centre sub-complex containing D1, D2, Cytochrome b559, PsbI and accessory factors
Chl a: Chlorophyll a
KFeCN: Potassium ferricyanide
DCBQ: 2,6-dichlorobenzoquinone.

