## Supplementary Information for "Cryo-EM structures of naturally occurring dimeric photosystem II complexes lacking the Mn_4_CaO_5_ cluster"

Zhao et al.

Supplementary Figures 1-27
Supplementary Tables 1- 4
Source data

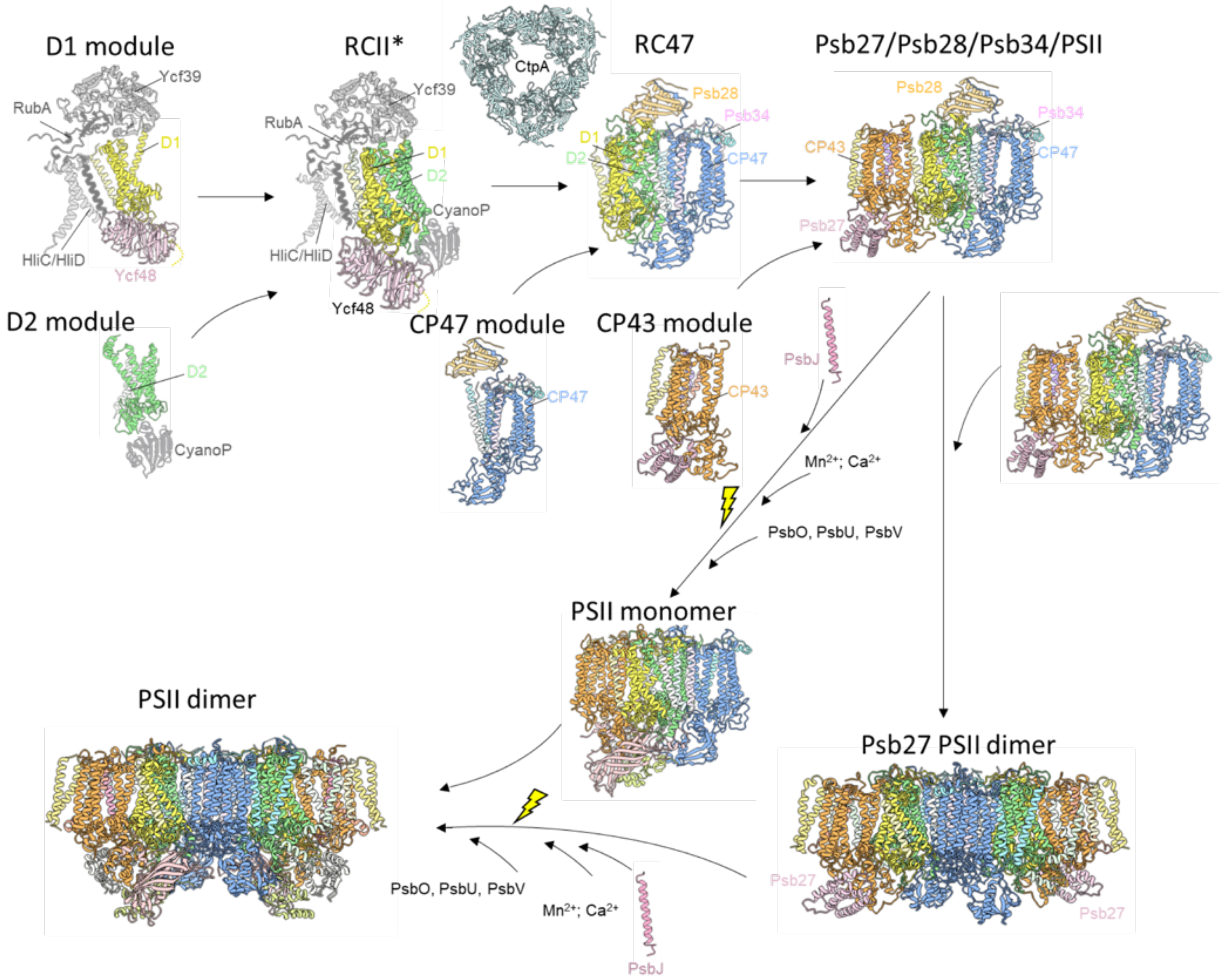


**Supplementary Fig 1. Model for assembly of PSII in cyanobacteria.**

PSII assembly begins with the formation of the D1 module, comprising precursor D1 (pD1), PsbI, Ycf48, RubA, and the Ycf39/Hlips complex. This module associates with the D2 module, which includes D2, cytochrome b-559 (Cyt b559), and CyanoP, to form the reaction center assembly complex (RCII*). The D1 C-terminal processing protease, CtpA, processes pD1. Subsequent addition of the CP47 module forms the RC47 complex, which is followed by the incorporation of the CP43 module, resulting in a non-oxygen-evolving PSII monomer. This intermediate then binds PsbJ and undergoes photoactivation along with attachment of the extrinsic subunits (PsbO, PsbU, and PsbV) to form a fully functional oxygen-evolving PSII monomer, which can further dimerize. Alternatively, PSII may first dimerize and then undergo photoactivation.

Key assembly factors and main PSII subunits are labelled. Protein structures are obtained from PDB with the ID in brackets: CtpA (8SXH); CyanoP (2XB3); D1 module; D2 module; RCII* (8ASL); CP47 module, CP43 module, RC47 and Psb27/Psb28/Psb34/PSII (7NHP); Psb27/PSII (7CZL); PSII monomer and PSII dimer (3WU2). Assembly cofactors not yet resolved in PSII complex are colored in grey. Remaining unresolved proteins are predicated by Alphafold (https://alphafoldserver.com/).

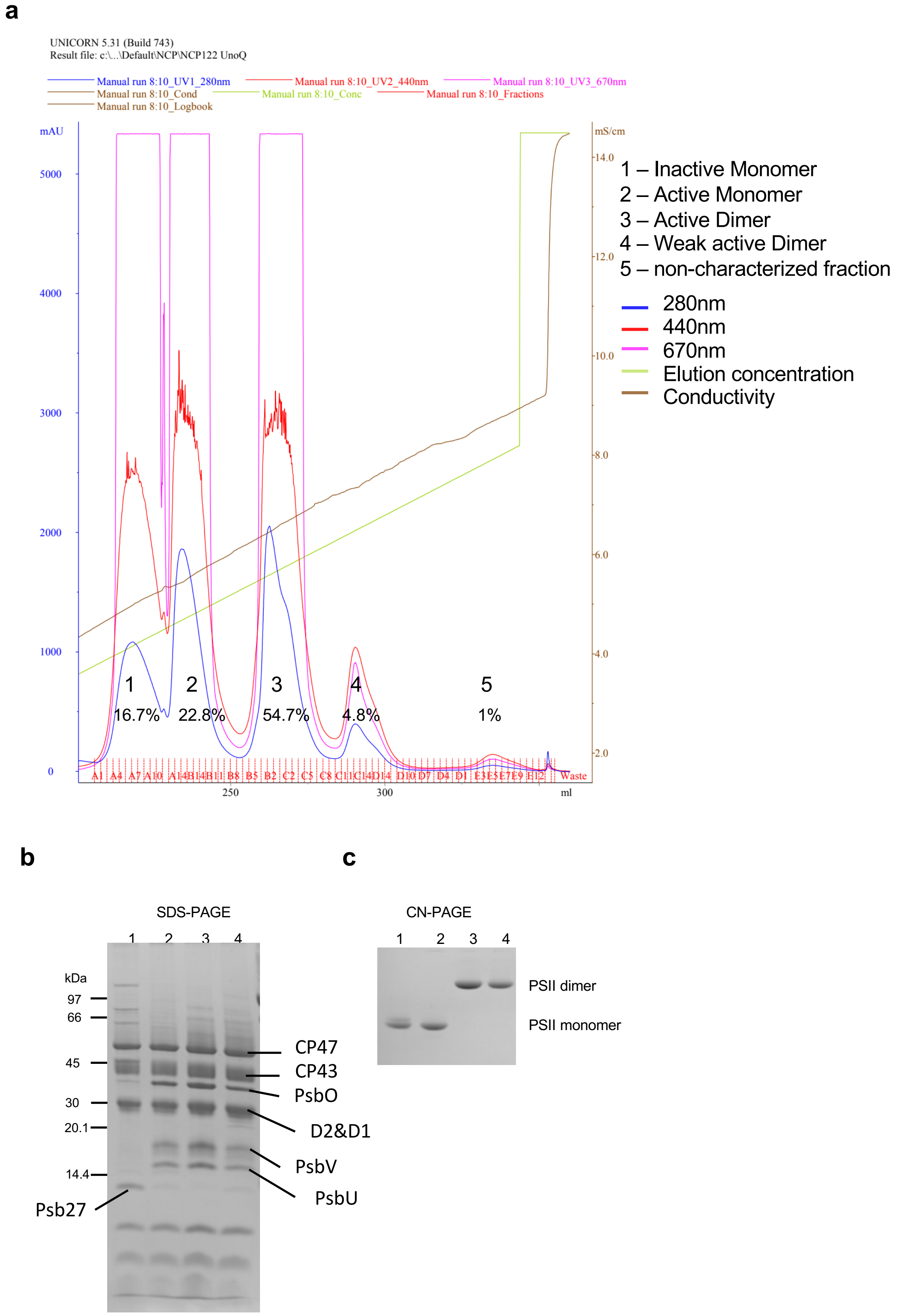


**Supplementary Fig 2. The anion exchange chromatography profile (a), the SDS-PAGE (b) and CN-PAGE (c) of four PSII fractions.**

**a.** Profile of anion exchange chromatography. PSII was eluted by 5-200 mM $MgSO_4$ gradient. Percentages of amount of PSII harvested are shown. Coomassie stained SDS-PAGE gel (**b**) and Clear Native-PAGE gel (**c**) of the four main peaks from anion exchange chromatography.

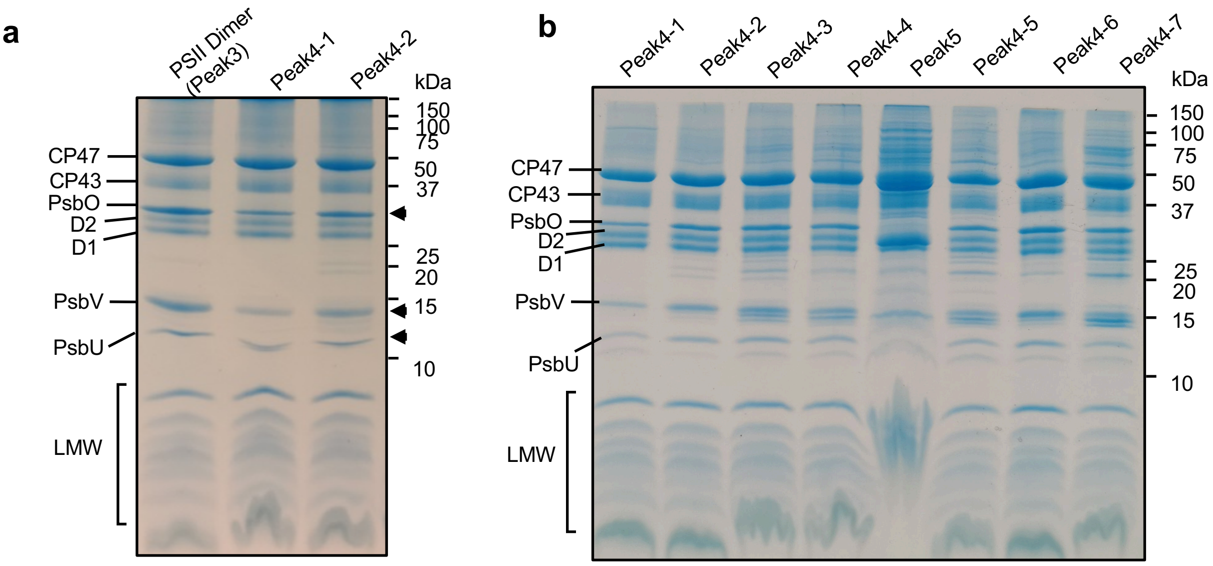


**c**

| Samples | Oxygen evolution Rate (μmol $O_2$ (mg Chl)$^{-1}$ $h^{-1}$) (Triplicates) |
|---|---|
| Active PSII Dimer | 3172 ± 344 |
| Peak4-1 | 1292 ± 300 |
| Peak4-2 | 2570 ± 293 |
| Peak4-3 | 1703 ± 251 |
| Peak4-4 | 2306 ± 441 |
| Peak5 | 837 ± NA |
| Peak4-5 | 1439 ± 543 |
| Peak4-6 | 3260 ± 1207 |
| Peak4-7 | 1307 ± 554 |

**Supplementary Fig 3. SDS-PAGE analysis and oxygen evolution activity of fully functional PSII (Peak3) and $PSII_{peak4}$. (a)** Coomassie stained SDS-PAGE gel of Peak3 and two samples of $PSII_{peak4}$ from two individual purifications. Equal amounts of chlorophyll were loaded for each sample. Peak4-2 was used for structural determination. **(b)** Coomassie stained SDS-PAGE gel of seven Peak4 samples (Peak4-1 to Peak4-7) from individual purifications and one Peak5 sample. "LMW" represents low-molecular-weight subunits. **(c)** Oxygen-evolution activity of Peak4 samples from **(b)**. Measurements were performed in triplicate.

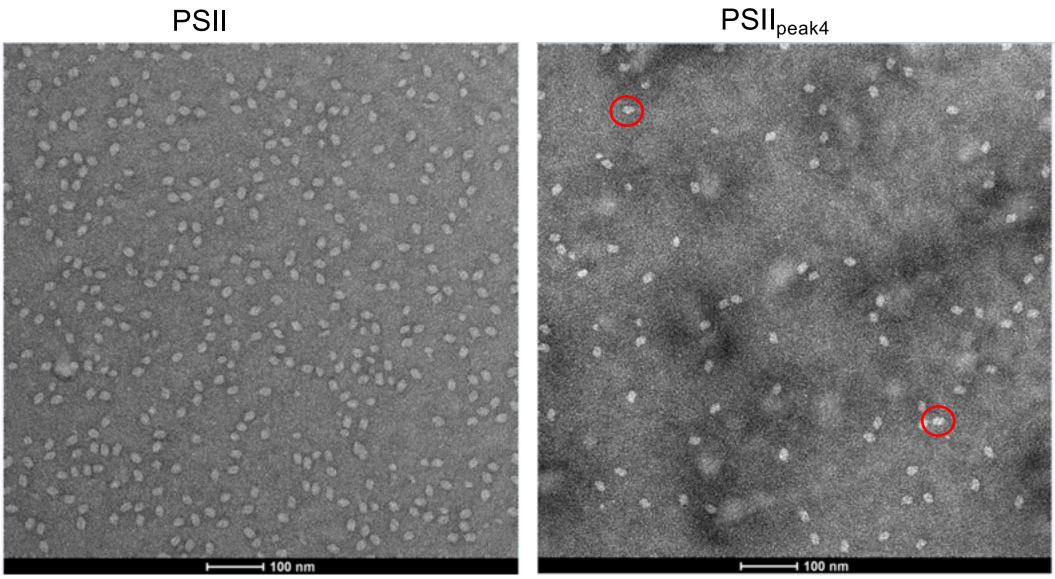


**Supplementary Fig 4. Negative staining of fully functional PSII and PSII$_{peak4}$.** Red circles highlight particles displaying dimeric features.

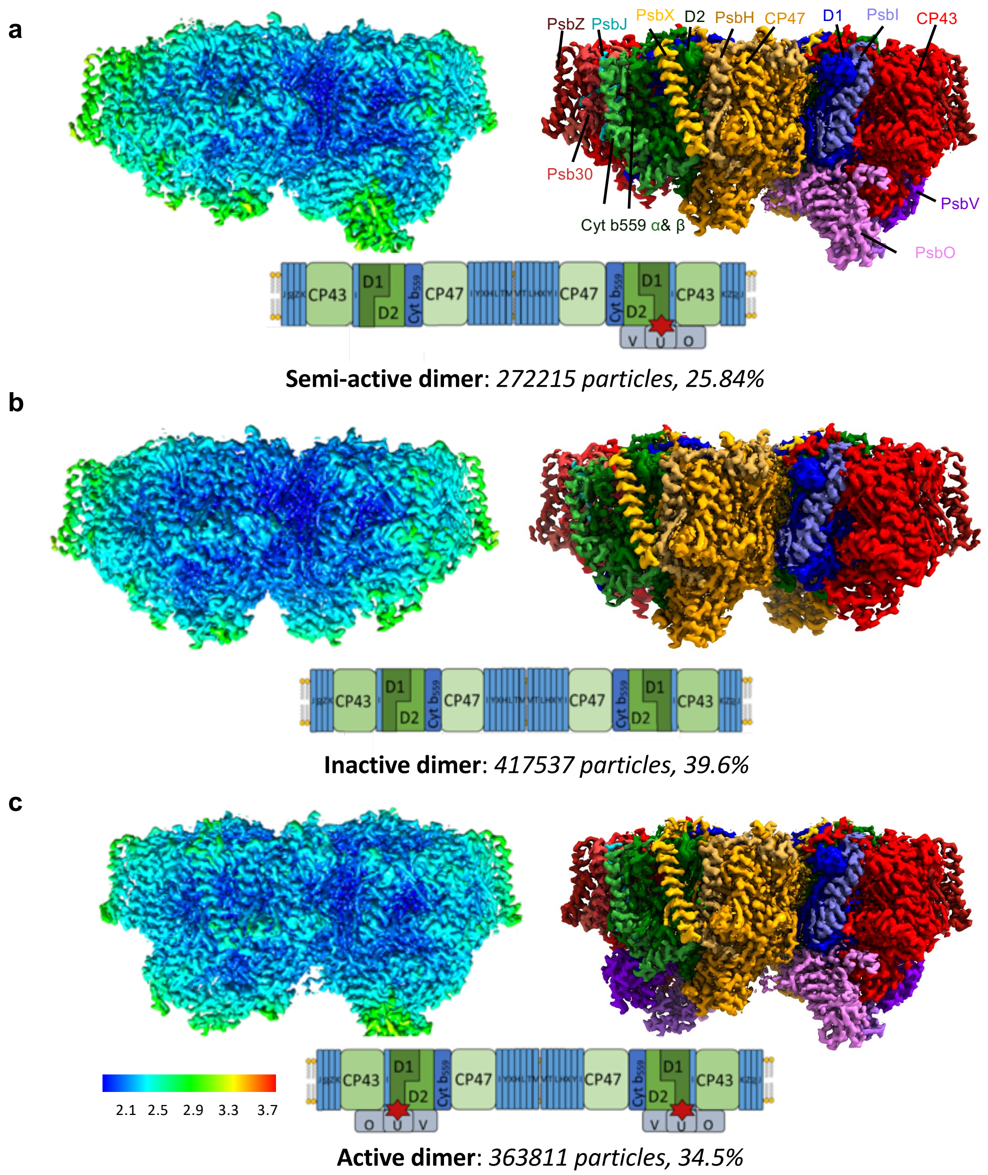


**Supplementary Fig 5. Local resolution of semi-active, inactive and active dimer maps.** **(a–c)** Cryo-EM maps for the semi-active **(a)**, inactive **(b)**, and active **(c)** PSII dimers. Left panels show maps colored by local resolution from high (blue) to low (red) resolution. Right panels show maps colored by subunit. Schematic cartoons below each map illustrate composition of each PSII complex. The percentages show proportion of particles contributing to each map out of total set of high-quality particles.

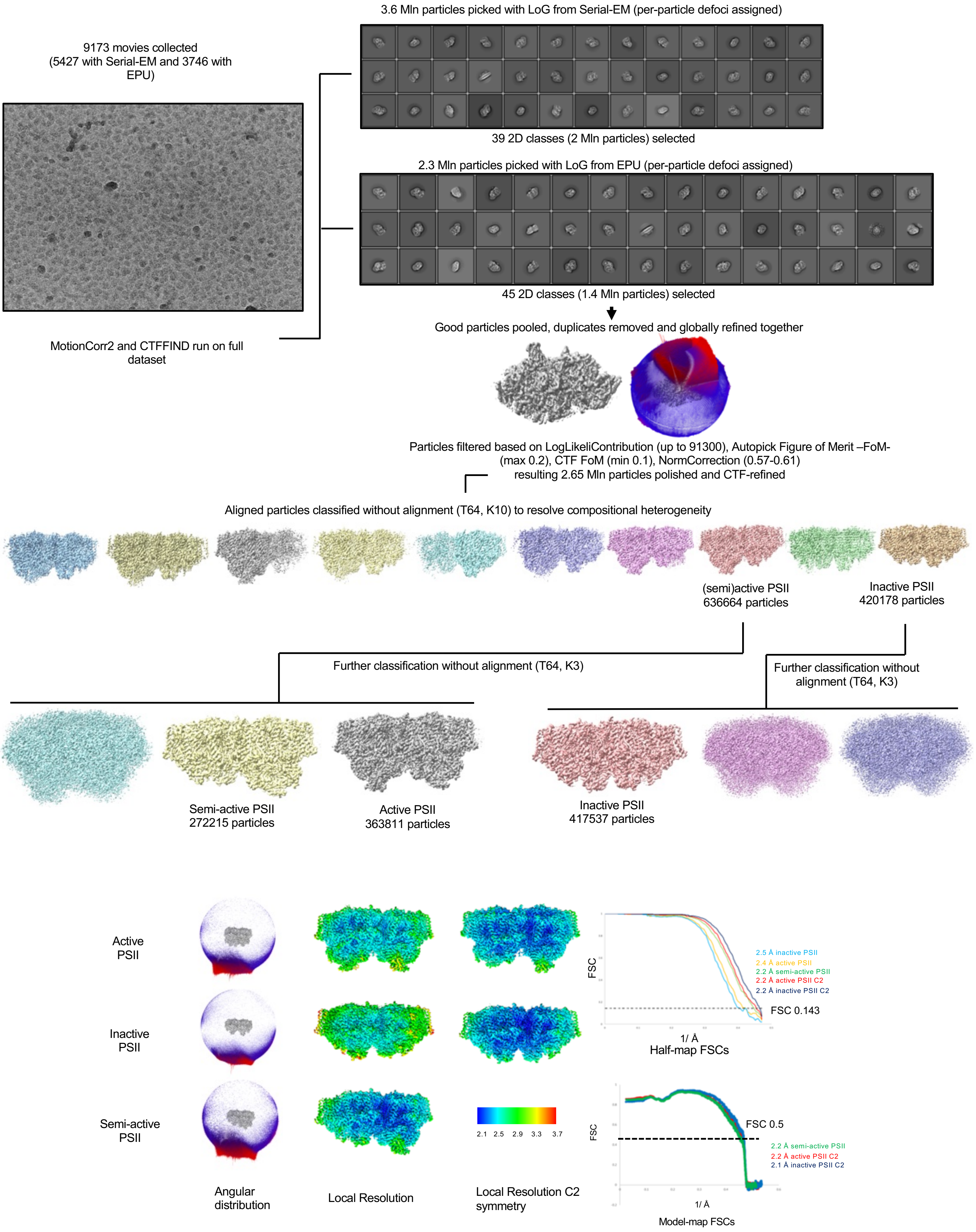


**Supplementary Fig 6. Data processing workflow.**

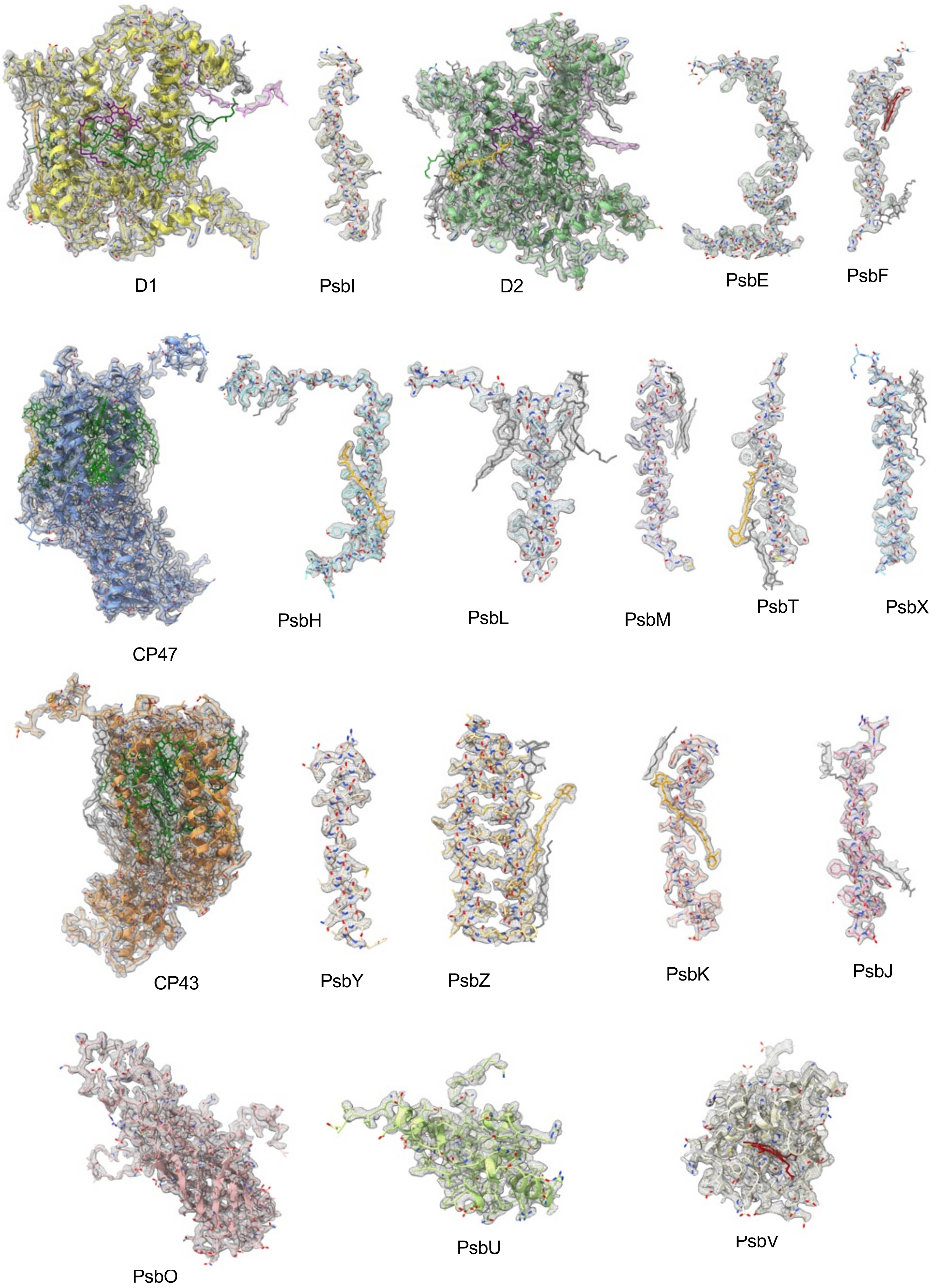


**Supplementary Fig 7. The density fit of active monomer subunits from the semi-active dimer.** Cryo-EM densities are shown at contour level 2σ. Cofactors are shown in different colors: Chlorophyll in green; carotenoids in orange; heme in maroon; other lipids/detergent in grey.

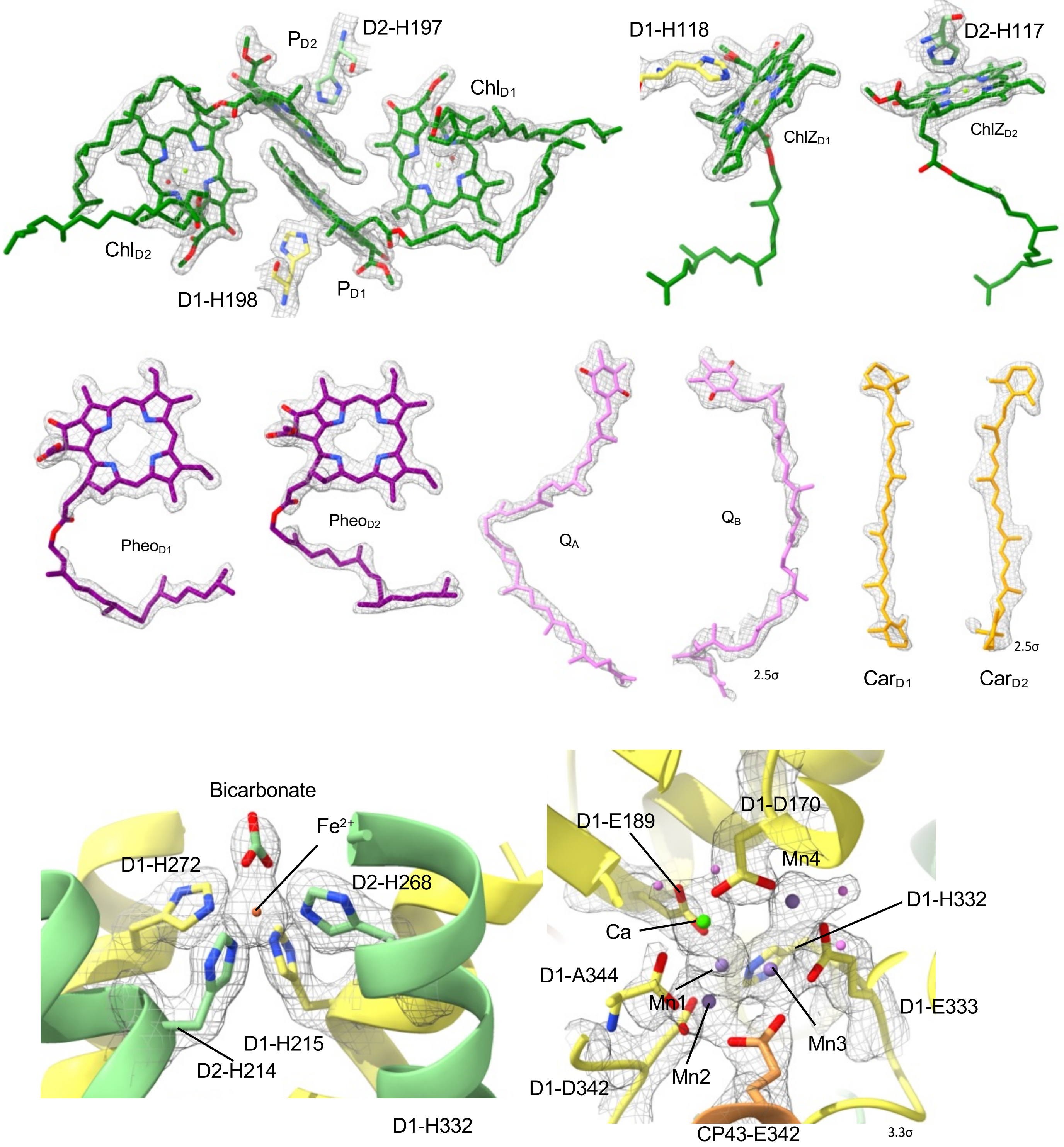


**Supplementary Fig 8. The density fit of cofactors from the active PSII monomer of the semi-active dimer.** The densities are shown at contour level 4σ, apart from $Q_B$ (2.5σ), $Car_{D2}$ (2.5σ) and the $Mn_4CaO_5$ (3.3σ) cluster.

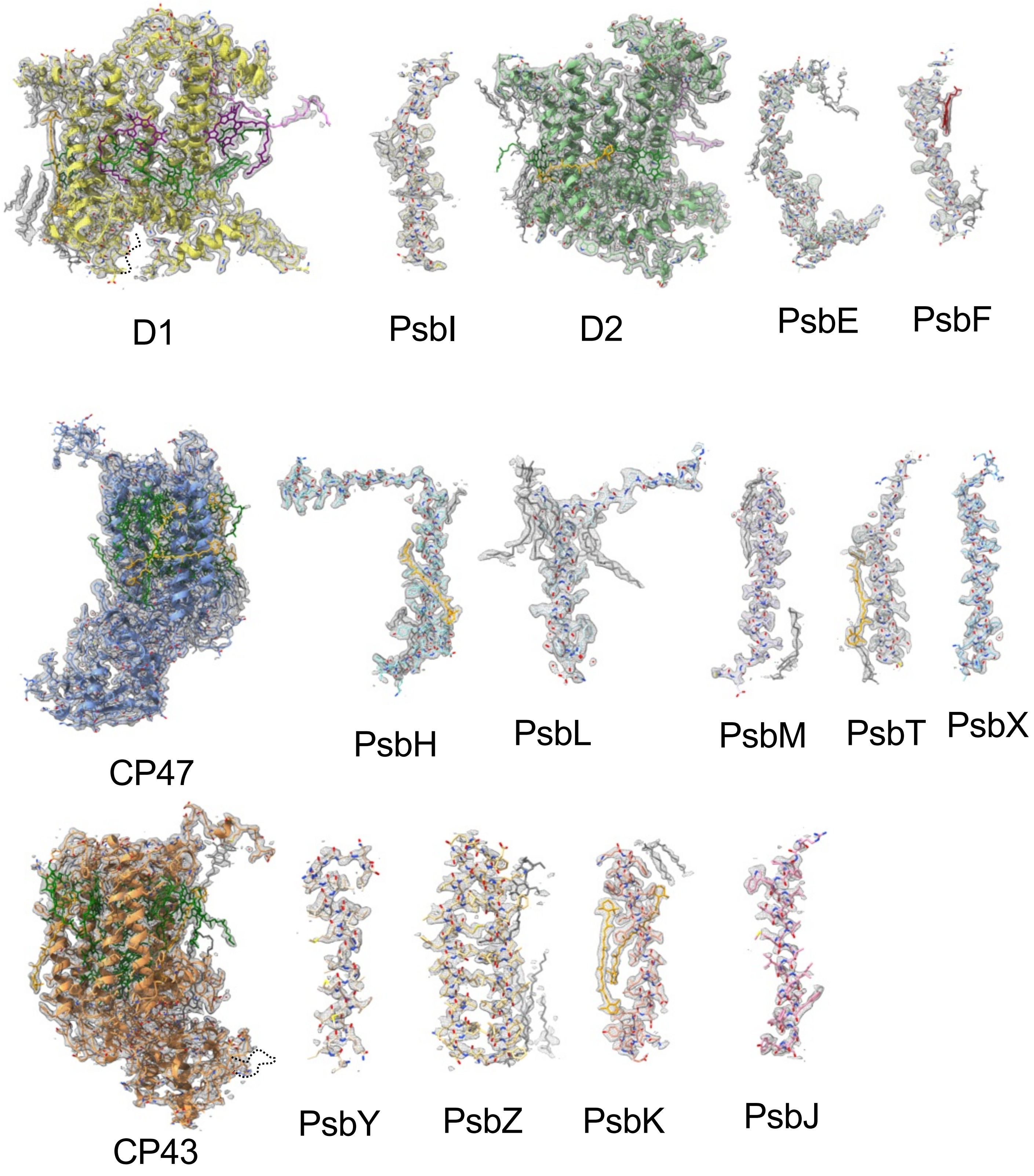


**Supplementary Fig 9. The density fit of inactive monomer subunits from the semi-active dimer.** Cryo-EM densities are shown at contour level 2 σ.

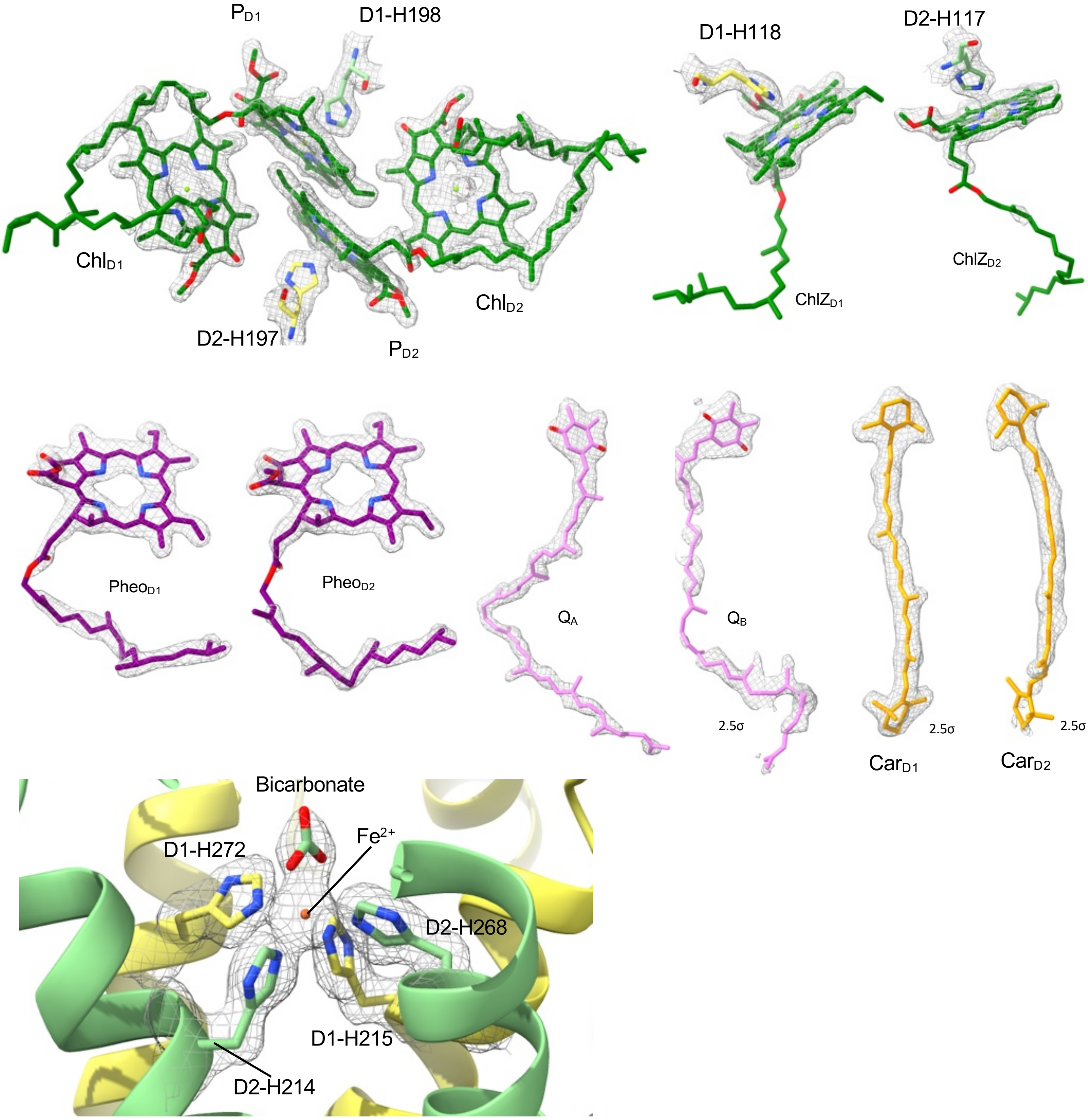


**Supplementary Fig 10. The density fit of key cofactors from the inactive PSII monomer from the semi-active dimer.** The densities are shown at contour level 4σ, apart from $Q_B$ (2.5σ), $Car_{D1}$ (2.5σ) and $Car_{D2}$ (2.5σ).

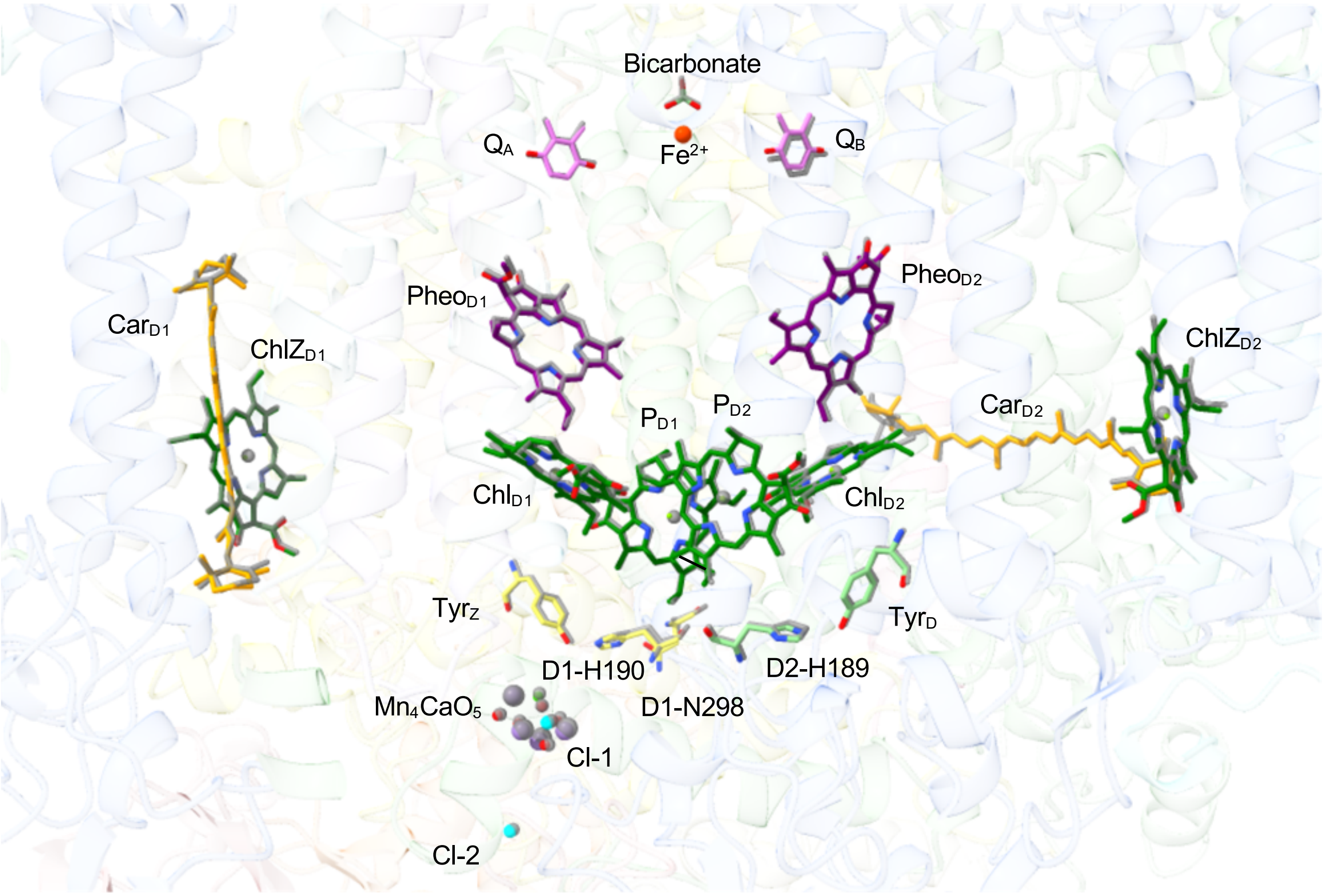


**Supplementary Fig 11. Distribution of key cofactors in the active monomer from semi-active dimer.**

Active monomer of the semi-active dimer is superimposed with the fully functional PSII (PDB ID: 3WU2). The cofactors of fully functional PSII are shown in grey, and the cofactors from the active monomer are shown in color.

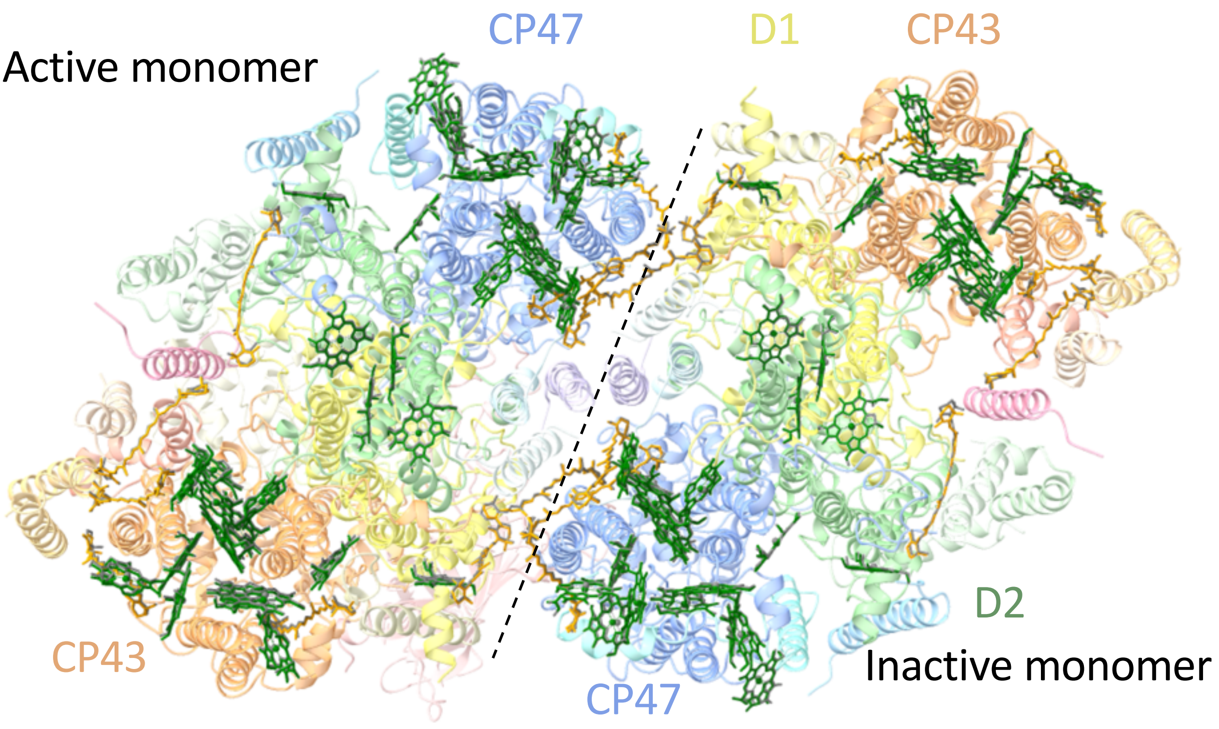


**Supplementary Fig 12. Chlorophyll and ß-carotene distribution in the semi-active dimer.**
All chlorophyll and ß-carotene molecules from fully functional PSII (PDB ID: 3WU2, grey) are present in the semi-active dimer (colored in green and orange). Chlorophyll tails are hidden for clarity.

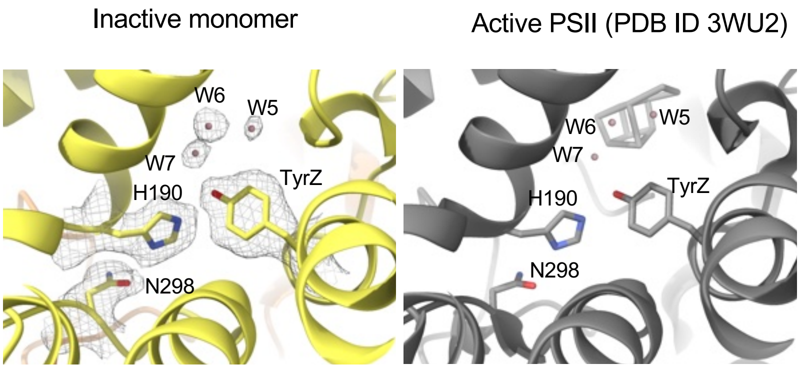


**Supplementary Fig. 13. Densities of water molecules around the $Y_Z$ in the inactive monomer.**

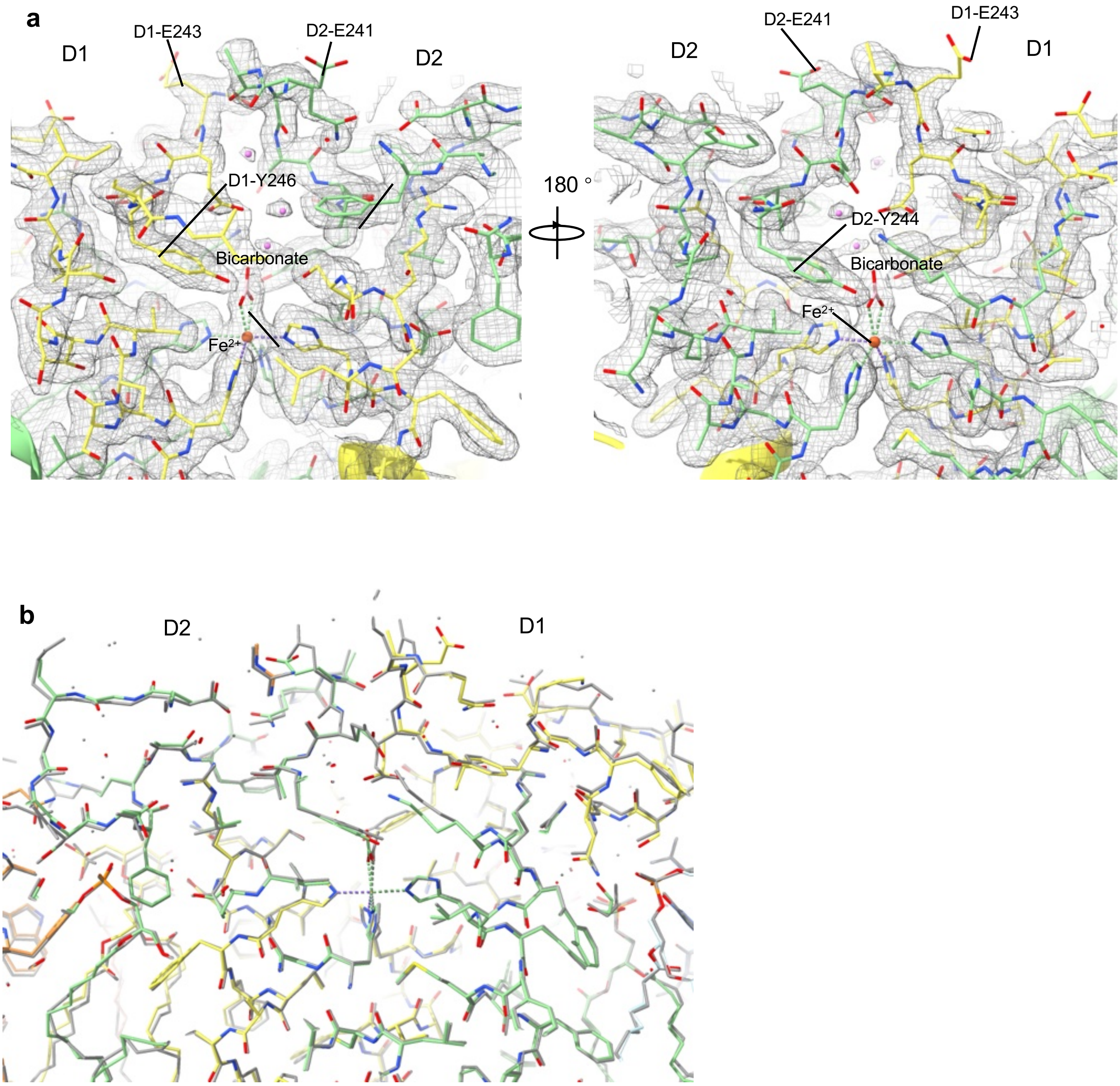


**Supplementary Fig. 14. Acceptor-side of inactive PSII monomer from semi-active PSII dimer.**
**a.** Cryo-EM density of acceptor-side of inactive PSII **b.** Superimposition of Inactive PSII (color) with fully functional PSII (PDB ID: 3WU2, grey).

**a**

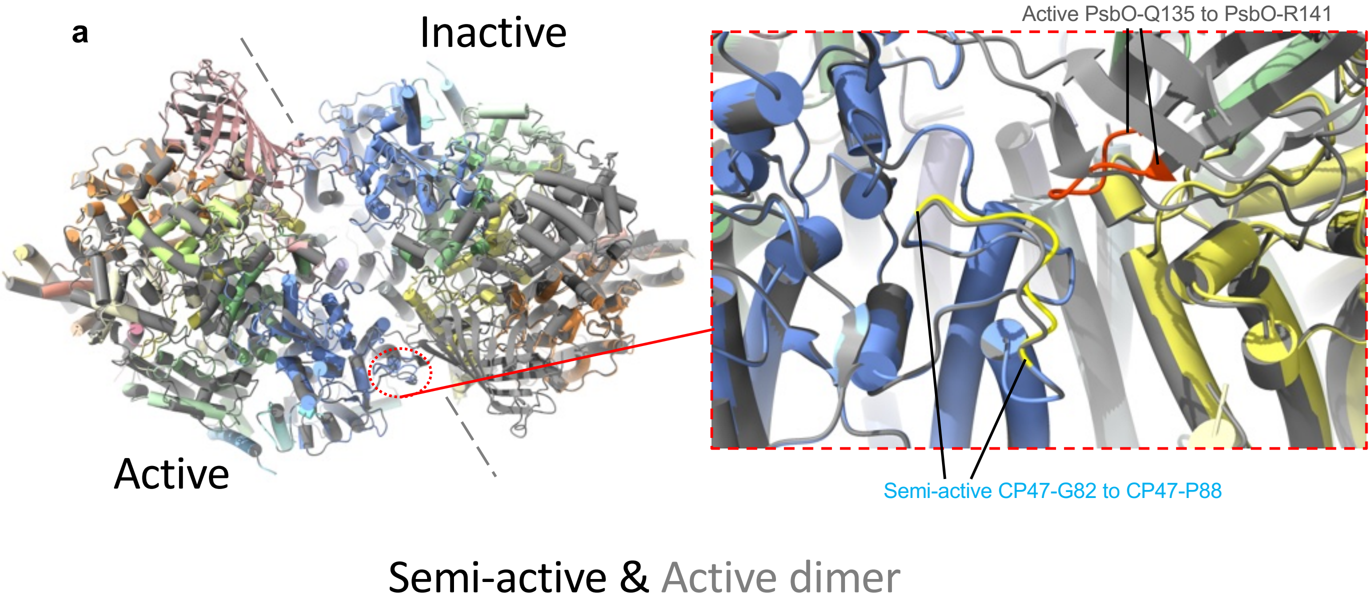


**b**

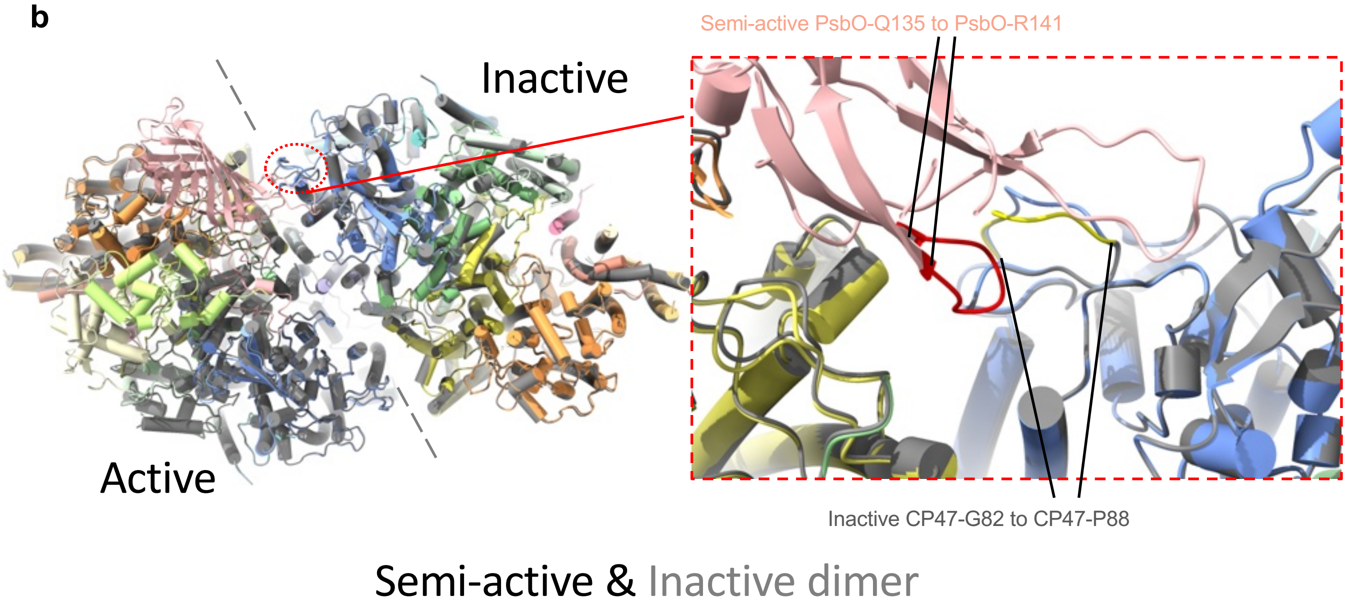


**Supplementary Fig. 15. Minor shift of the CP47 loop due to adjacent PsbO attachment.**

**a,** The semi-active dimer (colored) superimposed with the active dimer (grey). Luminal view (left) and close view of the monomer-monomer interface of PSII (right). The CP47 loop (yellow) of the active monomer (lacking adjacent PsbO) from the semi-active dimer clashes with the PsbO loop (red) from the active dimer. **b**, Superimposition of the semi-active PSII dimer (colored) with the inactive PSII dimer (grey), also shown from a luminal view (left) and close view (right). The PsbO loop (red) of the active monomer from the semi-active dimer clashes with the CP47 loop (yellow) from the inactive dimer.

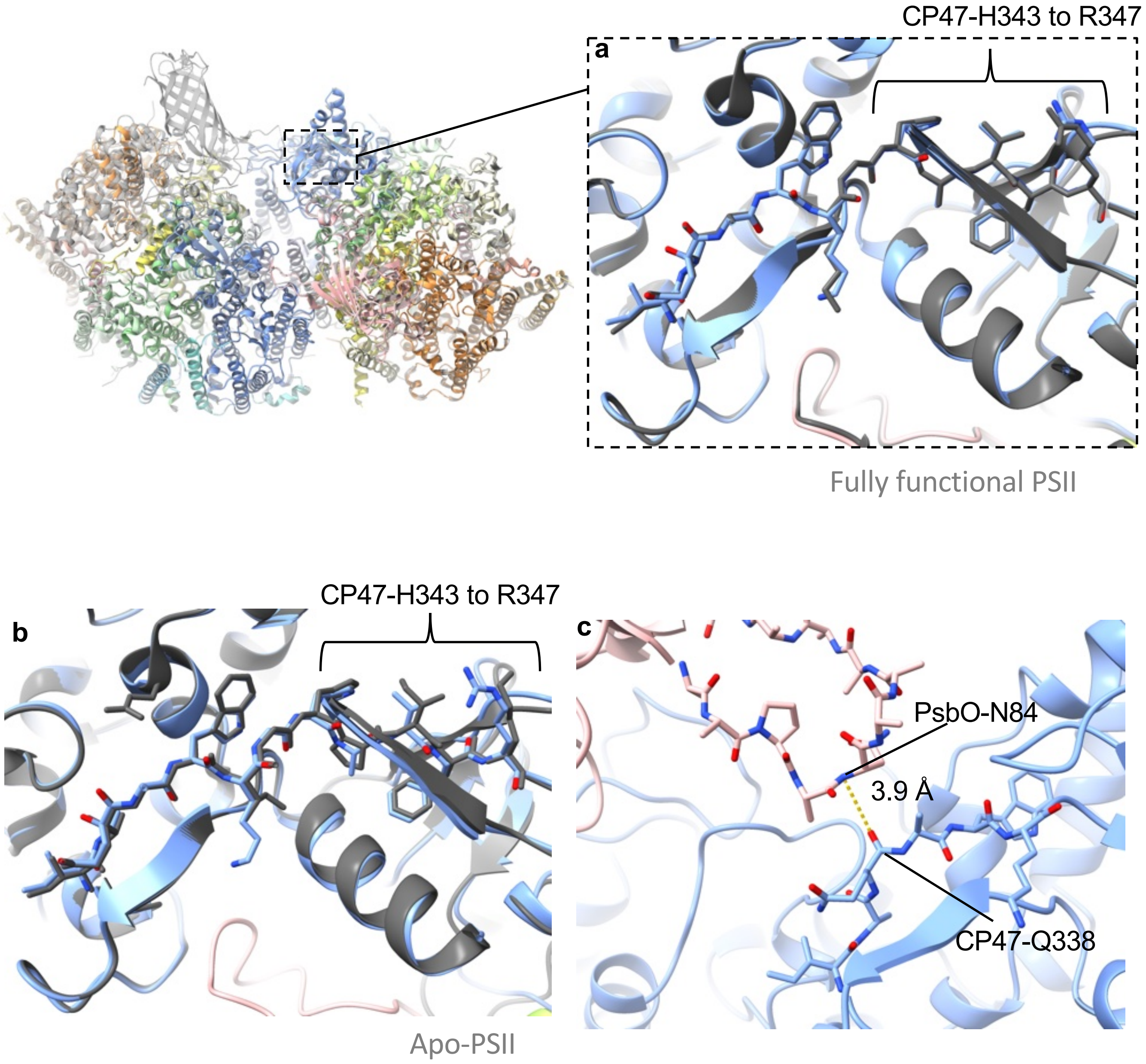


**Supplementary Fig. 16. Comparison between our active PSII monomer from the semi-active PSII dimer with fully functional PSII (a) and apo-PSII (b).**
**(a)** Active PSII monomer (blue; lacking adjacent PsbO) in the semi-active PSII dimer superimposed with fully functional PSII (PDB ID: 3wu2, grey). **(b)** Active PSII monomer (blue; lacking adjacent PsbO) in the semi-active PSII dimer superimposed with apo-PSII (6wj6, grey). **(c)** In the fully functional PSII (3wu2), at this region of ß sheet, only CP47-Q338 was found to have possible H-bond with the adjacent PsbO-N84 from the other PSII monomer.

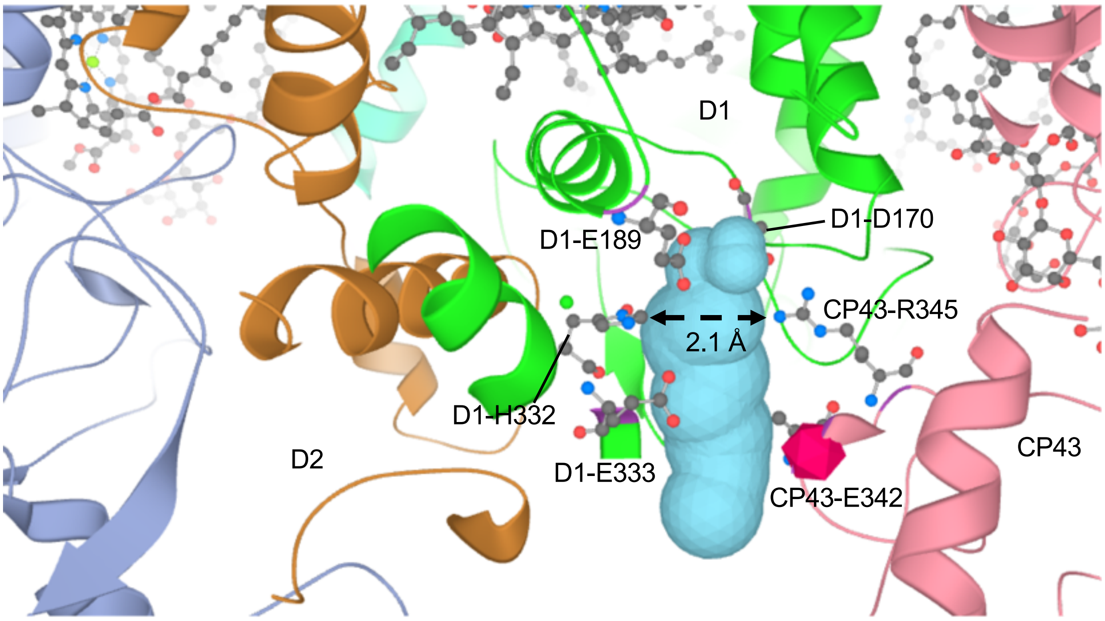


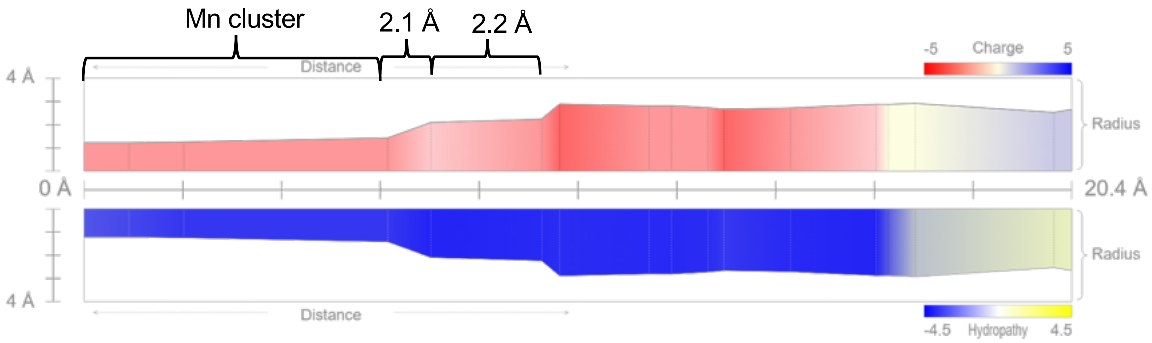


**Supplementary Fig. 17. The channel reaching the $Mn_4CaO_5$ cluster binding site in the inactive monomer from semi-active dimer.**

The channel is depicted as cyan surface (top panel). The inner radius, charge and hydropathy profile of the channel is shown at the bottom panel. The figure was made using MOLEonline: https://mole.upol.cz/

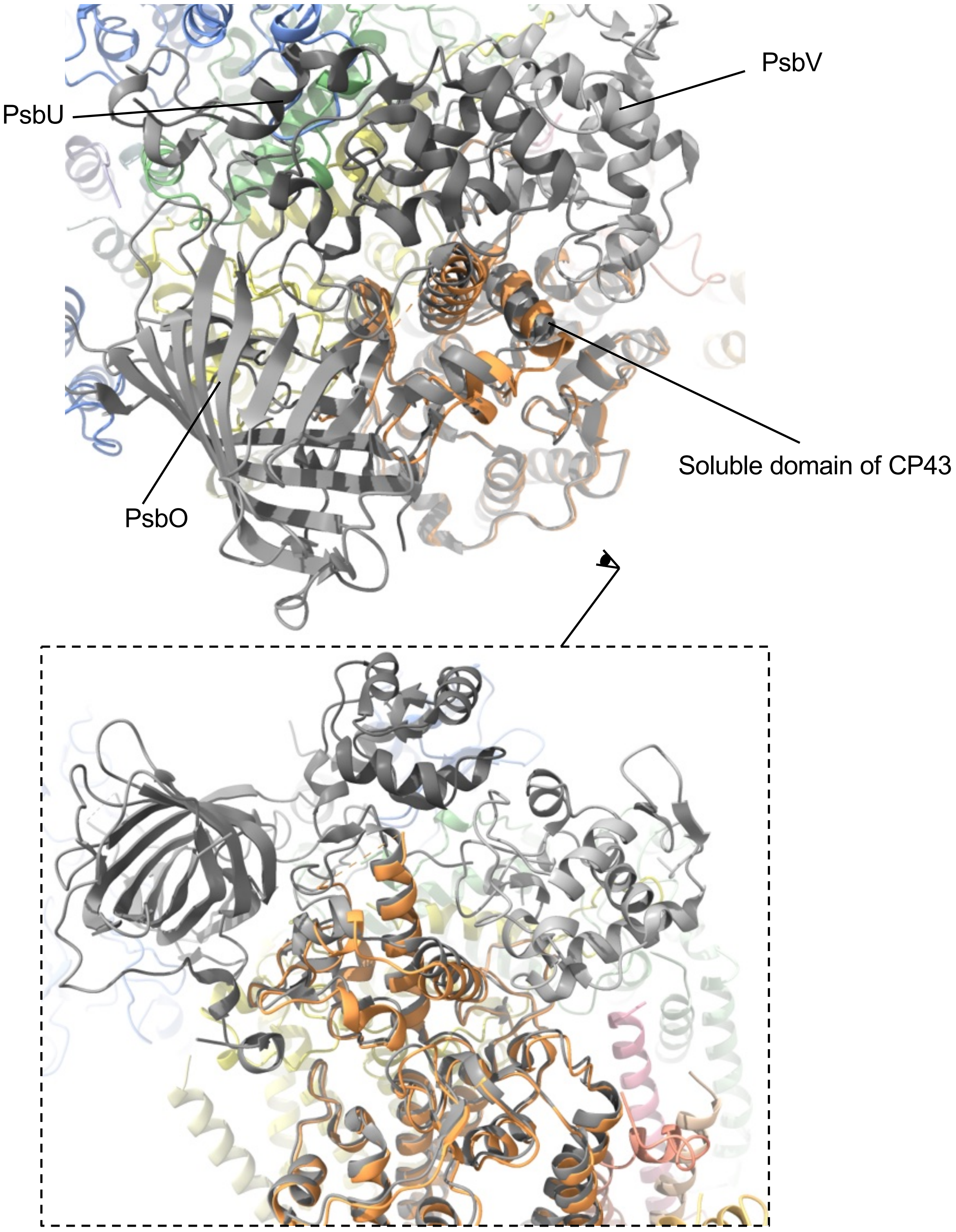


**Supplementary Fig. 18. Inactive monomer from the semi-active dimer superimposed with fully functional dimer.**
Inactive monomer is shown in color, CP43 is shown in orange. The fully functional PSII is shown in grey.

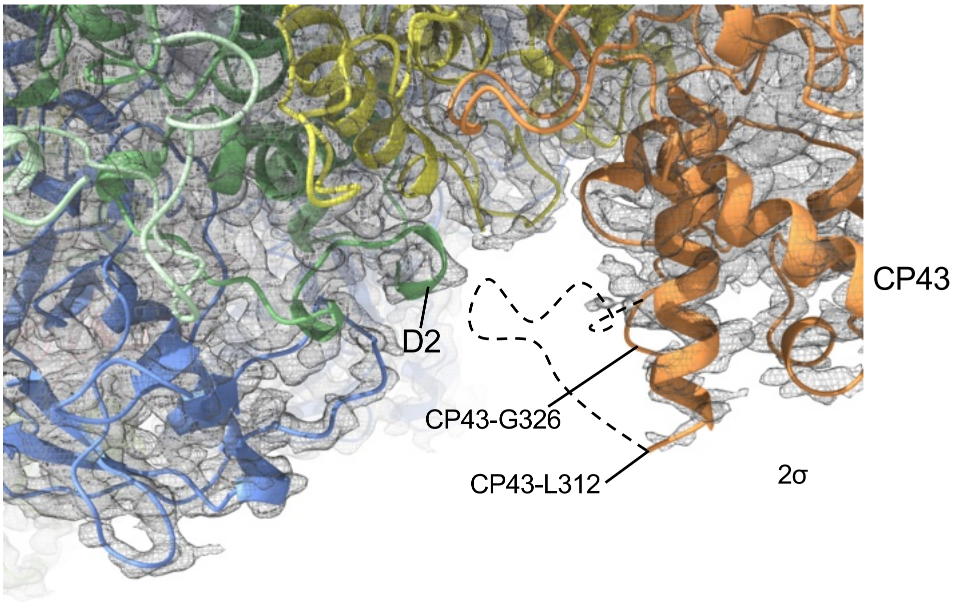


**Supplementary Fig. 19. Density fit of CP43 loop and D2 C-terminal at luminal side of PSII.**

Cryo-EM map is shown at contour level of 2σ.

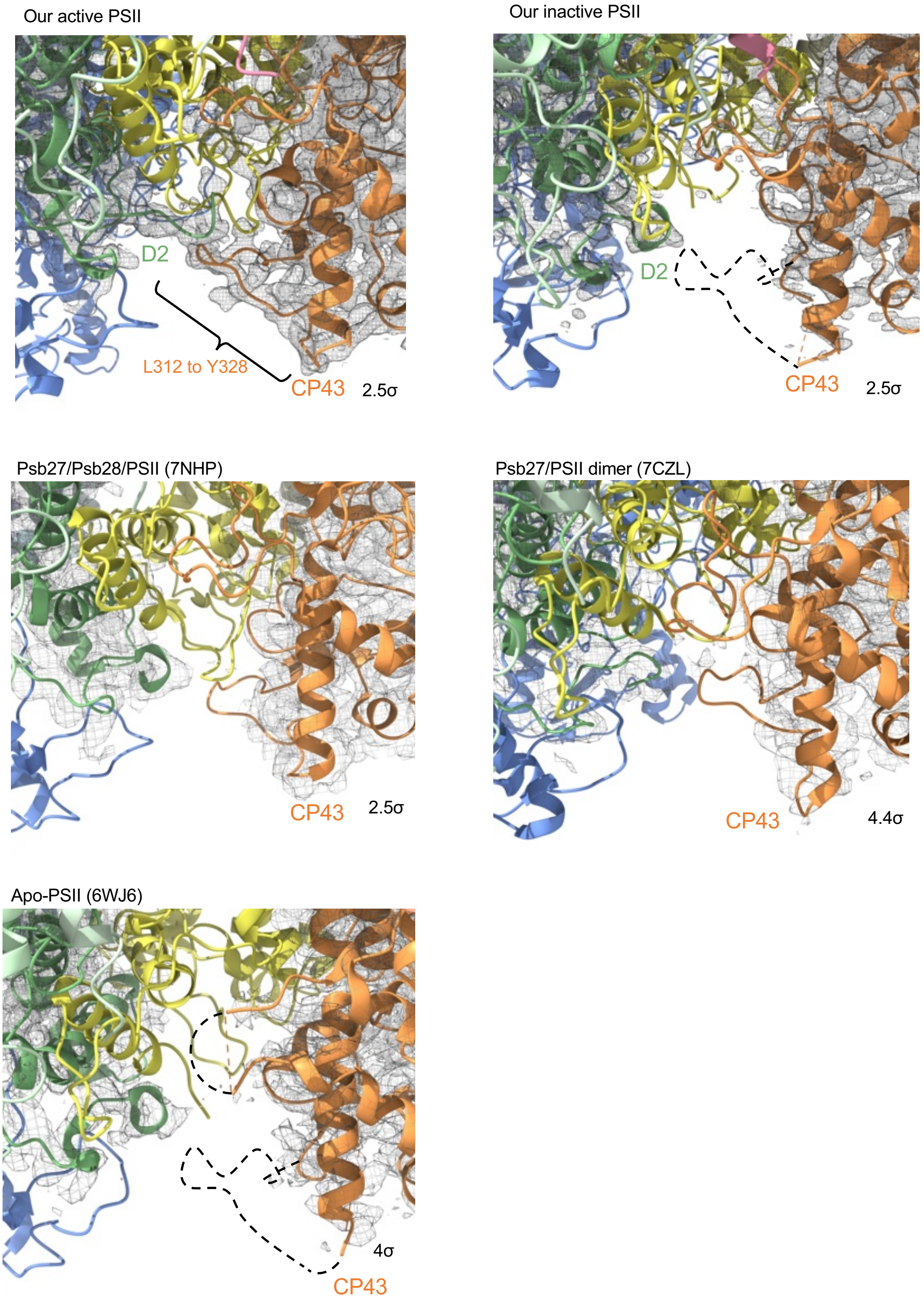


**Supplementary Fig. 20. Densities at the soluble domain of CP43 from L312 to Y328 of PSII intermediates.**

Extrinsic subunits and densities of D1 and CP47 were hidden for clarity. PDB IDs of structures are shown in brackets after the name of intermediates.

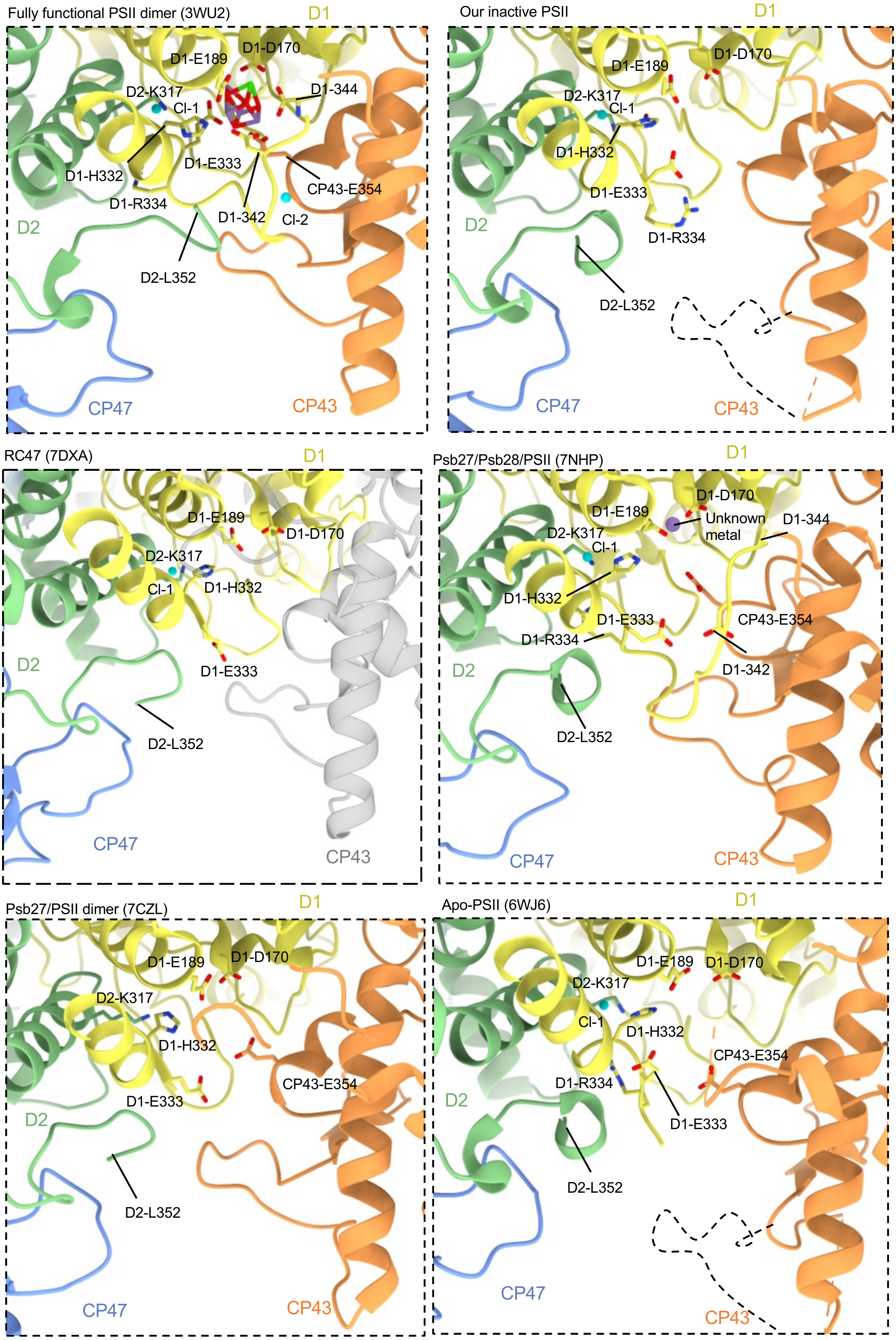


**Supplementary Fig. 21. Structural features at the donor side of PSII intermediates from our work and published structures.**
PDB ID of structures are shown in brackets after the name of intermediates.

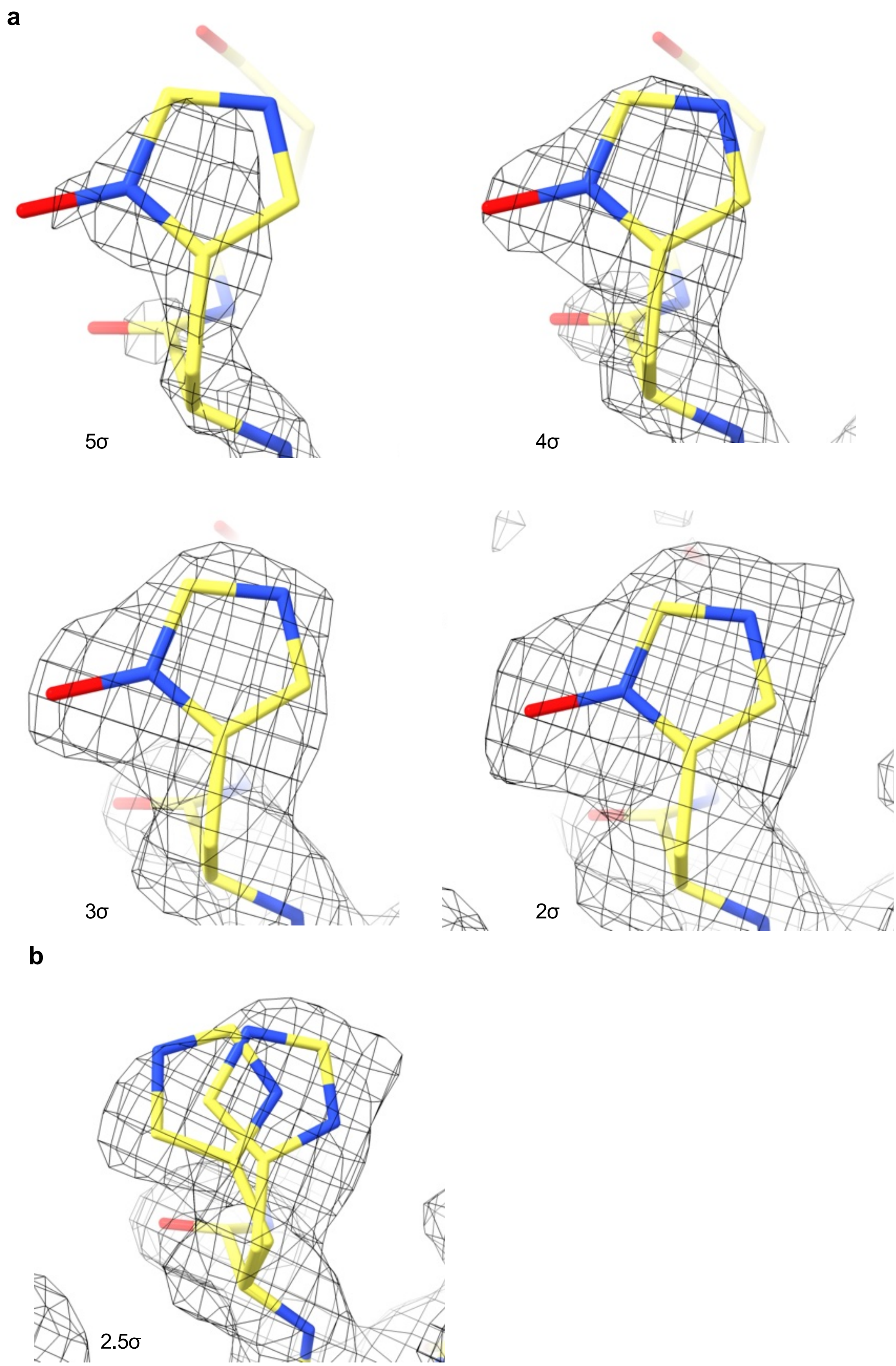


**Supplementary Fig. 22. D1-H332 density from inactive monomer of semi-active dimer.**
**(a)** Different contour levels of the inactive D1-H332. (**b)** Possible rotamers of histidine do not fit well into the density.

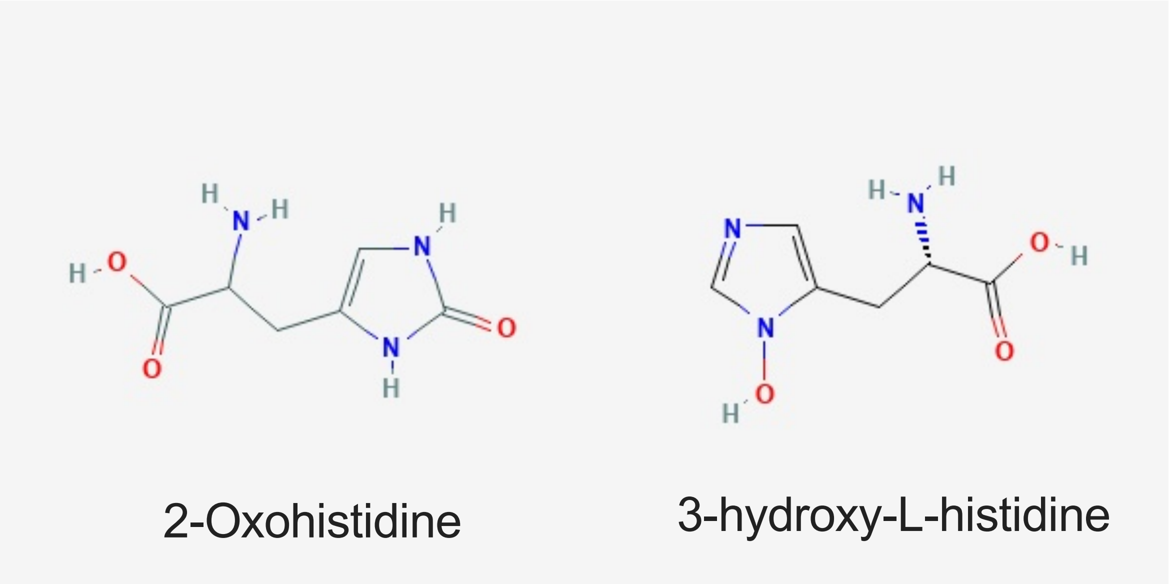


**Supplementary Fig. 23. 2-oxohistidine and hydroxy-histidine.**
Structures were taken from PDB.

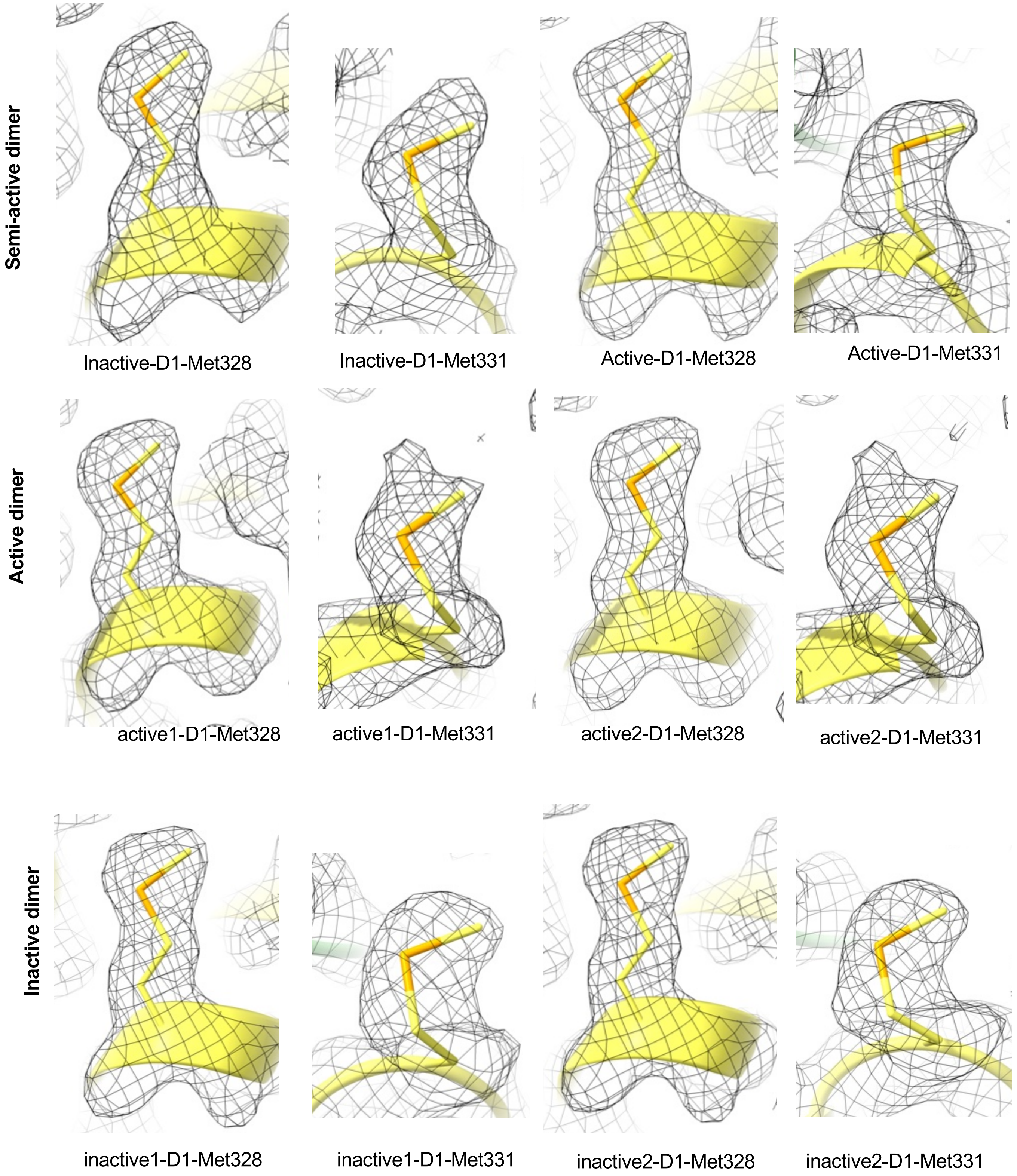


**Supplementary Fig. 24. The cryo-EM densities of methionine residues from D1 C-terminal tail.**
Densities are shown at contour level of 2σ.

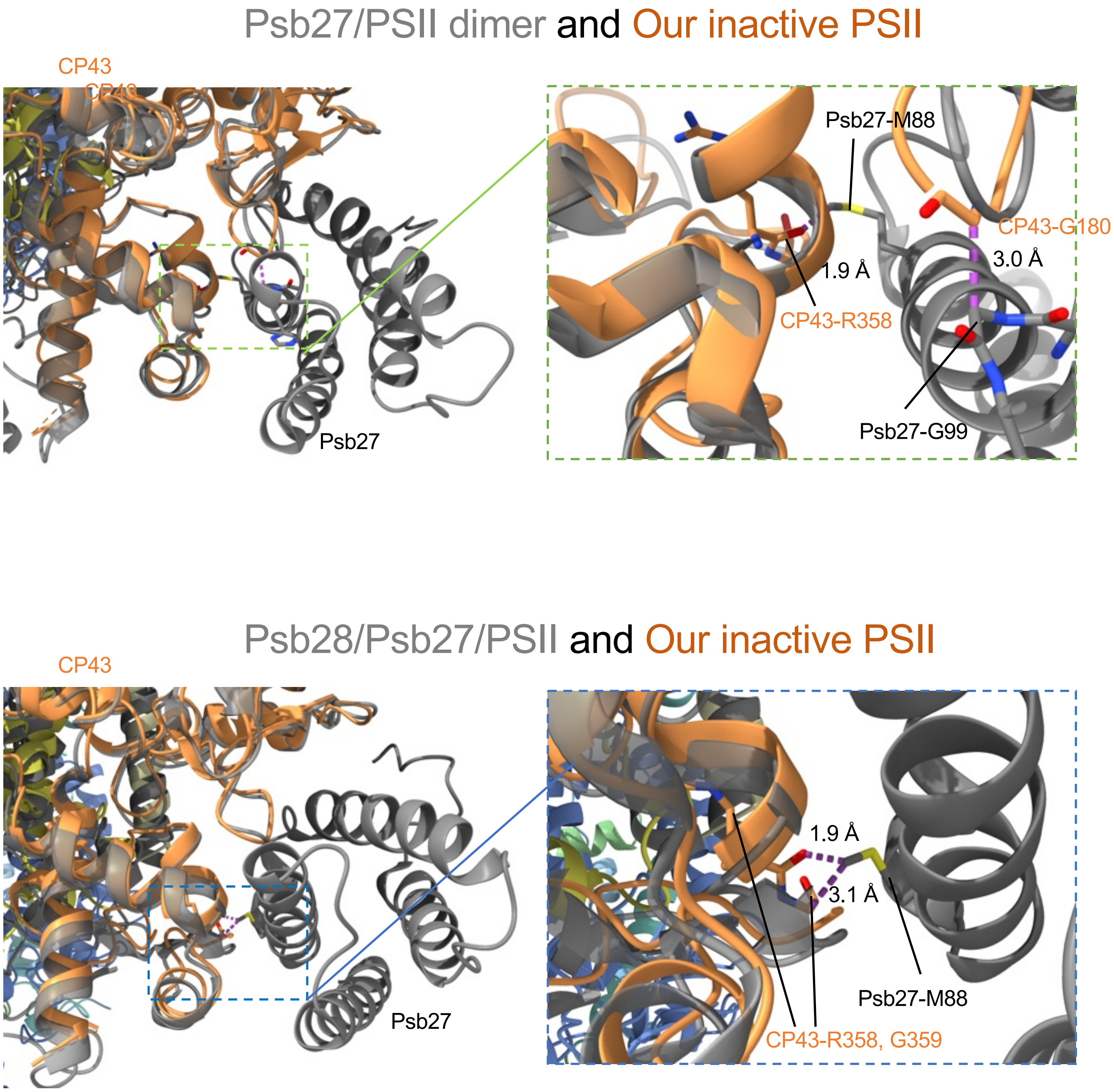


**Supplementary Fig. 25. CP43 of the inactive monomer was superimposed with the CP43 from the Psb27/PSII (PDB ID: 7CZL) dimer and the Psb28/Psb27/PSII (PDB ID: 7NHP).** The Psb27/PSII dimer and Psb28/Psb27/PSII are shown in grey and the inactive monomer subunits in color and labelled. The clashes are indicated with magenta dash lines. Clashes are identified in ChimeraX, when the sum of the atom radii of two atoms is 0.6 Å larger than the distance between the two atoms plus the potential H-bond allowance (0.4 Å).

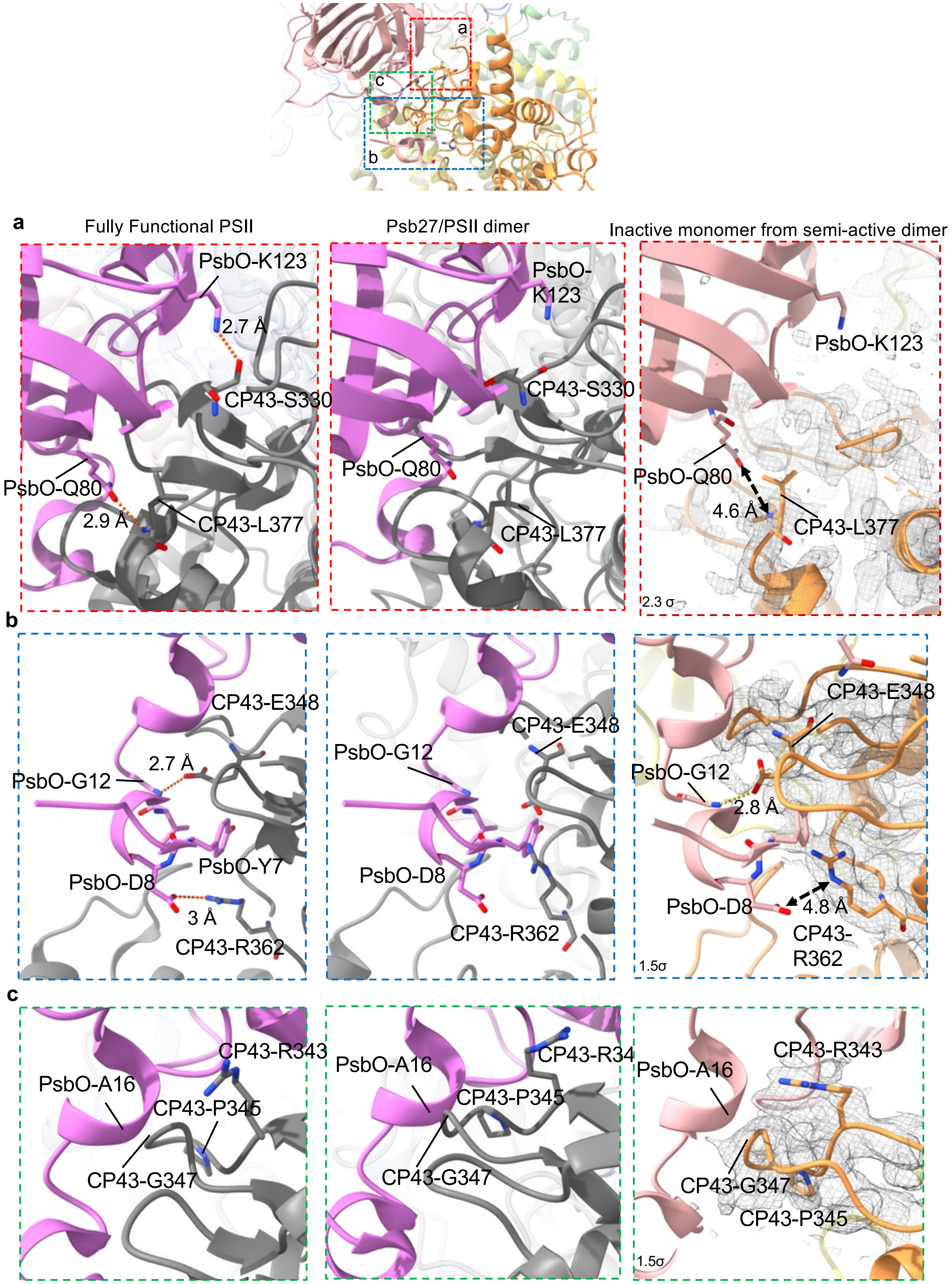


**Supplementary Fig. 26. CP43 and PsbO interactions in the fully functional PSII (PDB ID: 3WU2), Psb27/PSII ( PDB ID: 7CZL) and our Peak4 inactive monomer.** The PsbO is from the structure of fully functional PSII, which is superimposed with the inactive PSII monomer from the semi-active PSII dimer.

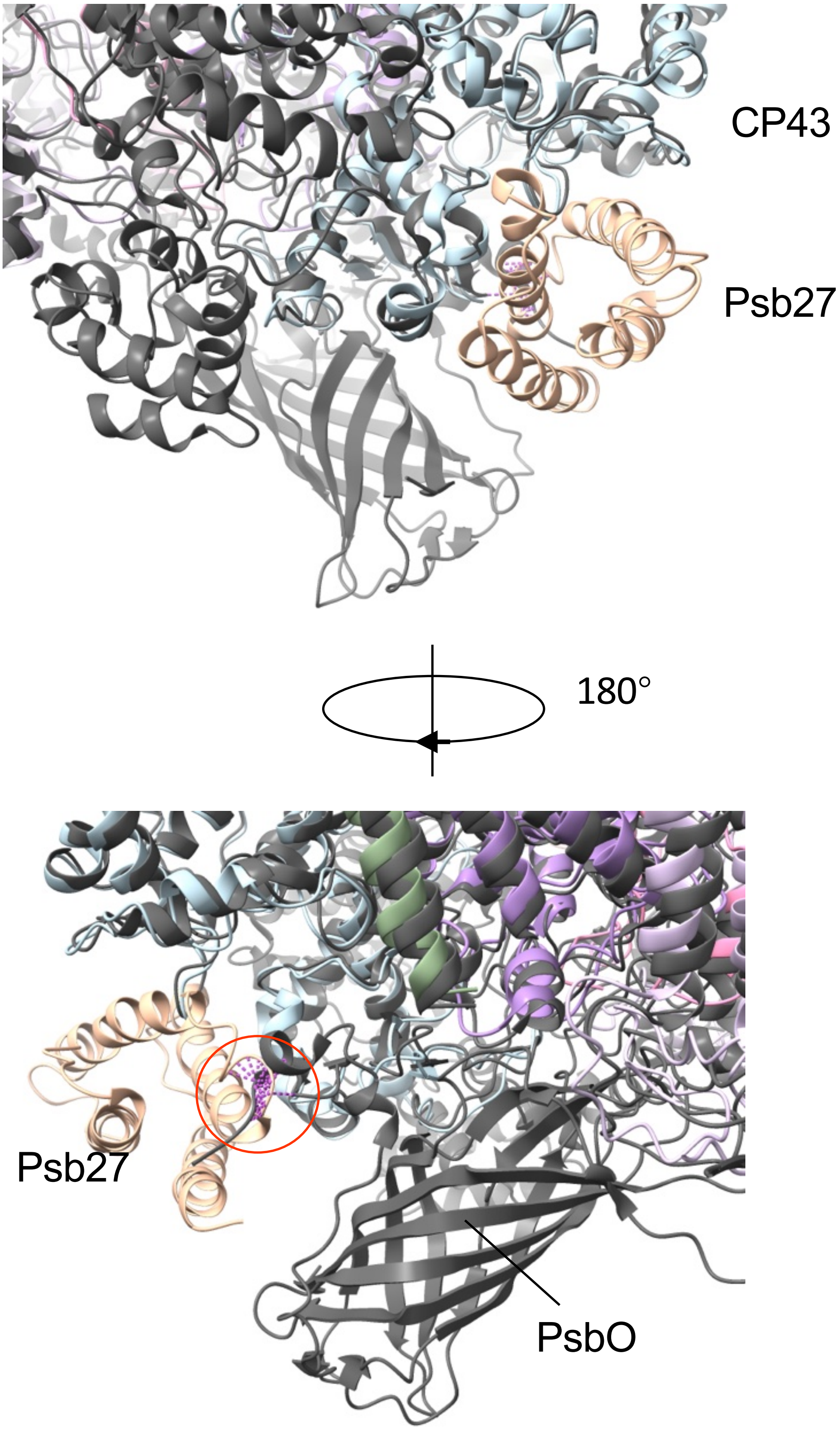


**Supplementary Fig. 27. CP43 soluble domain from Psb27/PSII (PDB ID: 7ZCL, colored) superimposed with that of fully functional PSII (PDB ID: 3WU2, grey).**
Clashes area is highlighted in magenta and red circle. Clashes are identified in ChimeraX, when the sum of the atom radii of two atoms is 0.6 Å larger than the distance between the two atoms plus the potential H-bond allowance (0.4 Å).

| | #1 Active PSII | #2 Inactive PSII | #3 Semi-active PSII |
|---|---|---|---|
| **Data collection and processing** | | | |
| Microscope | Titan Krios | Titan Krios | Titan Krios |
| Camera | K3 | K3 | K3 |
| Magnification | 81000X | 81000X | 81000X |
| Voltage (kV) | 300 | 300 | 300 |
| Electron exposure (e–/Å$^2$) | 80 | 80 | 80 |
| Automation software | Serial-EM/EPU | Serial-EM/EPU | Serial-EM/EPU |
| Number of frames | 80 | 80 | 80 |
| Defocus range (μm) | c.a. -0.6 to -2 | c.a. -0.6 to -2 | c.a. -0.6 to -2 |
| Pixel size (Å) | 1.061 | 1.061 | 1.061 |
| Tilt angle (degrees) | 30 | 30 | 30 |
| Symmetry imposed | C2 | C2 | C1 |
| Number of micrographs | 9173 | 9173 | 9173 |
| Initial particle images (no.) | 5.9M | 5.9M | 5.9M |
| Final particle images (no.) | 363811 | 417537 | 272215 |
| Map resolution (Å) at 0.143 FSC threshold | 2.21 | 2.15 | 2.23 |
| **Refinement** | | | |
| Initial model used (PDB code) | 7NHO (Inactive) | 3KZI (Active) | 7NHO (Inactive); 3KZI (Active) |
| Refinement package | Phenix_real _space_refinement | Phenix_real _space_refinement | Phenix_real _space_refinement |
| Model resolution (Å) at 0.5 FSC threshold | 2.22 | 2.15 | 2.24 |
| Local resolution range (Å) | 2.12 – 3.0 | 2.12 – 3.0 | 2.12 – 3.0 |
| Cross-correlation | | | |
| Mask | 0.90 | 0.91 | 0.89 |
| Volume | 0.88 | 0.89 | 0.88 |
| Map sharpening B factor (Å$^2$) | -28.8 | -29.5 | -24.0 |
| Model composition | | | |
| Non-hydrogen atoms | 51094 | 42661 | 46997 |
| Protein residues | 5239 | 4211 | 4727 |
| Ligands | 232 | 212 | 228 |
| Water | 1106 | 811 | 960 |
| B factors (Å$^2$) (mean) | | | |
| Protein | 39.49 | 44.95 | 48.41 |
| Ligand | 44.06 | 49.17 | 52.36 |
| Water | 35.98 | 39.82 | 41.91 |
| R.m.s. deviations | | | |
| Bond lengths (Å) | 0.004 | 0.005 | 0.005 |
| Bond angles (°) | 0.773 | 0.790 | 0.839 |
| Validation | | | |
| MolProbity score | 1.43 | 1.30 | 1.46 |
| Clashscore | 4.66 | 4.49 | 4.81 |
| Poor rotamers (%) | 1.78 | 1.24 | 1.89 |
| C-beta deviations | 0 | 0 | 0 |
| CaBLAM outliers (%) | 1.24 | 0.91 | 1.13 |
| Ramachandran plot | | | |
| Favored (%) | 98.53 | 98.50 | 98.30 |
| Allowed (%) | 1.47 | 1.45 | 1.70 |
| Disallowed (%) | 0.00 | 0.05 | 0.00 |

**Supplementary Table 1. Data processing statistics.**

| | Subunit name | Chain | Range built/total residues | Un-modelled residues | % modelled residues | Cofactors |
|---|---|---|---|---|---|---|
| Active monomer | D1 | A | 12-344/1-344 | 1-11 | 96.80 | chlorophyll, non-heme iron, β-carotene, pheophytin, $Mn_4CaO_5$ cluster, chloride, plastoquinone, SQD, LFA, PLM |
| | D2 | D | 12-352/1-352 | 1-11 | 96.88 | chlorophyll, pheophytin, β-carotene, bicarbonate, plastoquinone, LHG, LMG, DDM, DGD, SQD, LFA, PLM |
| | CP47 | B | 2-506/2-510 | 507-510 | 99.21 | chlorophyll, β-carotene, DGD, LMG, DDM, glycerol, LFA, PLM |
| | CP43 | C | 11-461/1-461 | 1-10 | 97.83 | chlorophyll, β-carotene, DGD, LMG, DDM, SQD, PLM, LFA |
| | PsbE | E | 4-83/2-84 | 2, 3, 84 | 96.39 | PLM, LFA |
| | PsbF | F | 11-45/2-45 | 2-10 | 79.55 | heme, DDM |
| | PsbH | H | 2-65/2-66 | 66 | 98.46 | RRX, LFA |
| | PsbI | I | 1-35/1-38 | 36-38 | 92.11 | LFA |
| | PsbJ | J | 6-40/2-40 | 2-5 | 89.74 | DDM, PLM |
| | PsbK | K | 10-46/10-46 | | 100.00 | β-carotene, PLM |
| | PsbL | L | 3-37/1-37 | 1-2 | 94.59 | LHG, SQD |
| | PsbM | M | 1-33/1-36 | 34-36 | 91.67 | DDM, PLM |
| | PsbT | T | 1-30/1-32 | 31,32 | 93.75 | β-carotene, DDM |
| | PsbX | X | 2-39/2-41 | 40,41 | 95.00 | SQD |
| | PsbY | Y | 16-46/1-46 | 1-15 | 67.39 | |
| | PsbZ | Z | 1-61/1-62 | 62 | 98.39 | LMG, DDM, β-carotene |
| | PsbO | O | 30-272/27-272 | 27-29 | 98.78 | Ca |
| | PsbU | U | 38-134/31-134 | 31-37 | 93.27 | |
| | PsbV | V | 27-163/27-163 | | 100.00 | heme |
| | Active monomer | | 2621/2712 | | 96.64 | |
| Inactive monomer | D1 | a | 12-334/1-344 | 1-11, 335-344 | 93.90 | chlorophyll, β-carotene, pheophytin, chloride, plastoquinone, SQD, LMG, DDM, PLM, LFA |
| | D2 | d | 12-352/1-352 | 1-11 | 96.88 | chlorophyll, non-heme iron, β-carotene bicarbonate, plastoquinone, DGD,SQD, LHG, LMG, LFA, PLM |
| | CP47 | b | 2-505/2-510 | 506-510 | 99.02 | chlorophyll, β-carotene, LMG, DDM, PLM, LFA |
| | CP43 | c | 12-312, 326-461/1-461 | 1-11, 313-325 | 94.79 | chlorophyll, β-carotene, DGD, LMG, DDM, PLM |
| | PsbE | e | 8-83/2-84 | 2-7, 84 | 91.57 | LHG, LFA |
| | PsbF | f | 12-45/2-45 | 2-11 | 77.27 | heme, DDM |
| | PsbH | h | 2-65/2-66 | 66 | 98.46 | RRX, DGD, PLM |
| | PsbI | i | 1-35/1-38 | 36-38 | 92.11 | LFA |
| | PsbJ | j | 6-34/2-40 | 2-5, 35-40 | 74.36 | |
| | PsbK | k | 10-46/10-46 | | 100.00 | β-carotene, PLM |
| | PsbL | l | 3-37/1-37 | 1-2 | 94.59 | LHG, SQD, LFA |
| | PsbM | m | 1-33/1-36 | 34-36 | 91.67 | DDM |
| | PsbT | t | 1-30/1-32 | 31, 32 | 93.75 | β-carotene, DDM |
| | PsbX | x | 2-39/2-41 | 40, 41 | 95.00 | |
| | PsbY | y | 18-46/1-46 | 1-17 | 63.04 | |
| | PsbZ | z | 1-61/1-62 | 62 | 98.39 | LMG, DDM, PLM |
| | Inactive monomer | | 2106/2225 | | 94.65 | |
| Total | Active/Inactive | | 4727/4937 | | 95.94 | |

SQD: Sulfoquinovosyl diacylglycerol; LHG: 1,2-dipalmitoyl-phosphatidyl-glycerole; LMG: 1,2-distearoyl-monogalactosyl-diglyceride; DDM: Dodecyl-beta-d-maltoside; DGD: Digalactosyl diacyl glycerol; RRX: (3R)-beta,beta-caroten-3-ol; PLM: Palmitic acid; LFA: Eicosane

**Supplementary Table 2. Statistics of modelling for the semi-active dimer.**

| | Subunit name | Chain | Range built/total residues | Un-modelled residues | % modelled residues | Cofactors |
|---|---|---|---|---|---|---|
| Active monomer 1 | D1 | a | 12-344/1-344 | 1-11 | 96.80 | chlorophyll, non-heme iron, β-carotene, pheophytin, $Mn_4CaO_5$ cluster, chloride, plastoquinone, SQD, DDM, LHG, PLM, LFA |
| | D2 | d | 12-352/1-352 | 1-11 | 96.88 | chlorophyll, pheophytin, β-carotene, bicarbonate, plastoquinone, LHG, LMG, DDM, DGD, PLM, LFA |
| | CP47 | b | 2-506/2-510 | 507-510 | 99.21 | chlorophyll, β-carotene, DGD, LMG, glycerol, DDM, PLM, LFA |
| | CP43 | c | 12-461/1-461 | 1-11 | 97.61 | chlorophyll, β-carotene, DGD, LMG, DDM, SQD, PLM, LFA |
| | PsbE | e | 4-84/2-84 | 2-3 | 97.59 | DDM, PLM |
| | PsbF | f | 12-45/2-45 | 2-11 | 77.27 | heme |
| | PsbH | h | 2-63/2-66 | 64-66 | 95.38 | RRX |
| | PsbI | i | 1-35/1-38 | 36-38 | 92.11 | LFA |
| | PsbJ | j | 5-40/2-40 | 2-4 | 92.31 | DDM, PLM |
| | PsbK | k | 10-46/10-46 | | 100.00 | β-carotene, PLM |
| | PsbL | l | 2-37/1-37 | 1 | 97.30 | LHG, SQD |
| | PsbM | m | 1-33/1-36 | 34-36 | 91.67 | DDM, PLM |
| | PsbT | t | 1-30/1-32 | 31-32 | 93.75 | β-carotene, DDM, PLM |
| | PsbX | x | 2-39/2-41 | 40-41 | 95.00 | SQD |
| | PsbY | y | 17-46/1-46 | 1-16 | 65.22 | |
| | PsbZ | z | 1-61/1-62 | 62 | 98.39 | LMG, DDM, β-carotene |
| | PsbO | o | 30-272/27-272 | 27-29 | 98.78 | Ca |
| | PsbU | u | 38-134/31-134 | 31-37 | 93.27 | |
| | PsbV | v | 27-163/27-163 | | 100.00 | heme |
| | Active monomer 1 | | 2619/2712 | | 96.57 | |
| Active monomer 2 | D1 | A | 12-344/1-344 | 1-11 | 96.80 | chlorophyll, non-heme iron, β-carotene, pheophytin, $Mn_4CaO_5$ cluster, chloride, plastoquinone, SQD, LHG, PLM, LFA |
| | D2 | D | 12-352/1-352 | 1-11 | 96.88 | chlorophyll, pheophytin, β-carotene, bicarbonate, plastoquinone, LHG, LMG, DDM, DGD, PLM |
| | CP47 | B | 2-506/2-510 | 507-510 | 99.21 | chlorophyll, β-carotene, DGD, LMG, glycerol, DDM, PLM, LFA |
| | CP43 | C | 12-461/1-461 | 1-11 | 97.61 | chlorophyll, β-carotene, DGD, LMG, DDM, SQD, PLM, LFA |
| | PsbE | E | 4-84/2-84 | 2, 3 | 97.59 | PLM |
| | PsbF | F | 12-45/2-45 | 2-11 | 77.27 | heme, PLM |
| | PsbH | H | 2-63/2-66 | 64-66 | 95.38 | RRX, PLM, LFA |
| | PsbI | I | 1-36/1-38 | 37-38 | 94.74 | LFA |
| | PsbJ | J | 5-40/2-40 | 2-4 | 92.31 | DDM , LFA |
| | PsbK | K | 10-46/10-46 | | 100.00 | β-carotene |
| | PsbL | L | 2-37/1-37 | 1 | 97.30 | LHG, SQD |
| | PsbM | M | 1-33/1-36 | 34-36 | 91.67 | DDM, PLM |
| | PsbT | T | 1-30/1-32 | 31, 32 | 93.75 | β-carotene, DDM, LFA |
| | PsbX | X | 2-39/2-41 | 40, 41 | 95.00 | SQD |
| | PsbY | Y | 17-46/1-46 | 1-16 | 65.22 | |
| | PsbZ | Z | 1-61/1-62 | 62 | 98.39 | LMG, DDM, β-carotene |
| | PsbO | O | 30-272/27-272 | 27-29 | 98.78 | Ca |
| | PsbU | U | 38-134/31-134 | 31-37 | 93.27 | |
| | PsbV | V | 27-163/27-163 | | 100.00 | heme |
| | Active monomer 2 | | 2620/2712 | | 96.61 | |
| Total | Active/Active | | 5239/5424 | | 96.59 | |

SQD: Sulfoquinovosyl diacylglycerol; LHG: 1,2-dipalmitoyl-phosphatidyl-glycerole; LMG: 1,2-distearoyl-monogalactosyl-diglyceride; DDM: Dodecyl-beta-d-maltoside; DGD: Digalactosyl diacyl glycerol; RRX: (3R)-beta,beta-caroten-3-ol; PLM: Palmitic acid; LFA: Eicosane

**Supplementary Table 3 Statistics of modelling for the active dimer.**

| | Subunit name | Chain | Range built/total residues | Un-modelled residues | % modelled residues | Cofactors |
|---|---|---|---|---|---|---|
| Inactive monomer 1 | D1 | A | 11-334/1-344 | 1-10, 335-344 | 94.19 | chlorophyll, non-heme iron, β-carotene, pheophytin, chloride, plastoquinone, SQD, LMG, DDM, PLM, LFA |
| | D2 | D | 12-352/1-352 | 1-11 | 96.88 | chlorophyll, β-carotene bicarbonate, plastoquinone, LHG, LMG, DGD, LFA |
| | CP47 | B | 2-505/2-510 | 506-510 | 99.02 | chlorophyll, β-carotene, LMG, DDM, PLM, LFA |
| | CP43 | C | 11-461/1-461 | 1-10 | 97.83 | chlorophyll, β-carotene, DGD, LMG, PLM, LFA |
| | PsbE | E | 4-84/2-84 | 2, 3 | 97.59 | PLM |
| | PsbF | F | 13-45/2-45 | 2-12 | 75.00 | heme, SQD |
| | PsbH | H | 2-65/2-66 | 66 | 98.46 | RRX, DGD, PLM, LFA |
| | PsbI | I | 1-35/1-38 | 36-38 | 92.11 | PLM |
| | PsbJ | J | 7-34/2-40 | 2-6, 35-40 | 71.79 | LFA |
| | PsbK | K | 10-46/10-46 | | 100.00 | β-carotene, LFA |
| | PsbL | L | 3-37/1-37 | 1, 2 | 94.59 | LHG, SQD |
| | PsbM | M | 1-33/1-36 | 34-36 | 91.67 | DDM, LFA |
| | PsbT | T | 1-30/1-32 | 31, 32 | 93.75 | β-carotene, DDM |
| | PsbX | X | 2-36/2-41 | 37-41 | 87.50 | |
| | PsbY | Y | 19-46/1-46 | 1-18 | 60.87 | |
| | PsbZ | Z | 1-61/1-62 | 62 | 98.39 | LFA |
| | Inactive monomer 1 | | 2120/2225 | | 95.28 | |
| Inactive monomer 2 | D1 | a | 11-334/1-344 | 1-10, 335-344 | 94.19 | chlorophyll, non-heme iron, β-carotene, pheophytin, chloride, plastoquinone, SQD, LMG, PLM, LFA |
| | D2 | b | 12-352/1-352 | 1-11 | 96.88 | chlorophyll, β-carotene bicarbonate, plastoquinone, LHG, LMG, DDM, DGD, LFA |
| | CP47 | c | 2-506/2-510 | 507-510 | 99.21 | chlorophyll, β-carotene, LMG, PLM, LFA |
| | CP43 | d | 11-461/1-461 | 1-10 | 97.83 | chlorophyll, β-carotene, DGD, DDM, LMG |
| | PsbE | e | 4-84/2-84 | 2, 3 | 97.59 | LHG, PLM |
| | PsbF | f | 13-45/2-45 | 2-12 | 75.00 | heme, SQD |
| | PsbH | h | 2-65/2-66 | 66 | 98.46 | RRX, DGD, LFA |
| | PsbI | i | 1-35/1-38 | 36-38 | 92.11 | PLM, LFA |
| | PsbJ | j | 7-34/2-40 | 2-6, 35-40 | 71.79 | |
| | PsbK | k | 10-46/10-46 | | 100.00 | β-carotene, LFA |
| | PsbL | l | 3-37/1-37 | 1, 2 | 94.59 | LHG, SQD |
| | PsbM | m | 1-33/1-36 | 34-36 | 91.67 | DDM, LFA |
| | PsbT | t | 1-30/1-32 | 31, 32 | 93.75 | β-carotene, DDM |
| | PsbX | x | 2-36/2-41 | 37-41 | 87.50 | |
| | PsbY | y | 19-46/1-46 | 1-18 | 60.87 | |
| | PsbZ | z | 1-61/1-62 | 62 | 98.39 | DDM |
| | Inactive monomer 2 | | 2121/2225 | | 95.33 | |
| Total | Inactive/Inactive | | 4241/4450 | | 95.30 | |

SQD: Sulfoquinovosyl diacylglycerol; LHG: 1,2-Dipalmitoyl-phosphatidyl-glycerole; LMG: 1,2-Distearoyl-monogalactosyl-diglyceride; DDM: Dodecyl-beta-d-maltoside; DGD: Digalactosyl diacyl glycerol; RRX: β-Cryptoxanthin; HTG: Heptyl 1-thio-beta-D-glucopyranoside

**Supplementary Table 4. Statistics of modelling for the inactive dimer.**

### Source Data

b

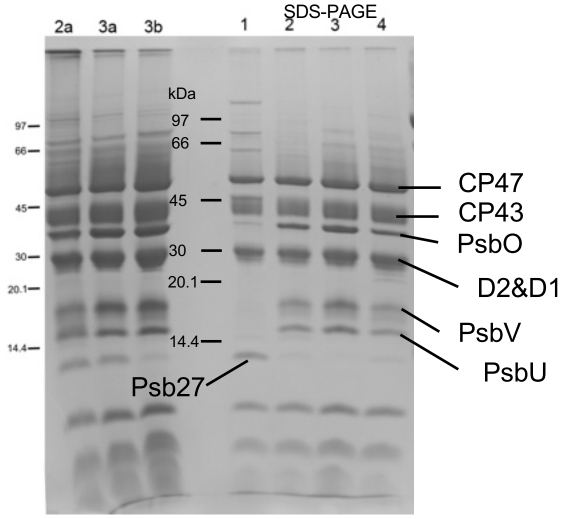


c CN-PAGE

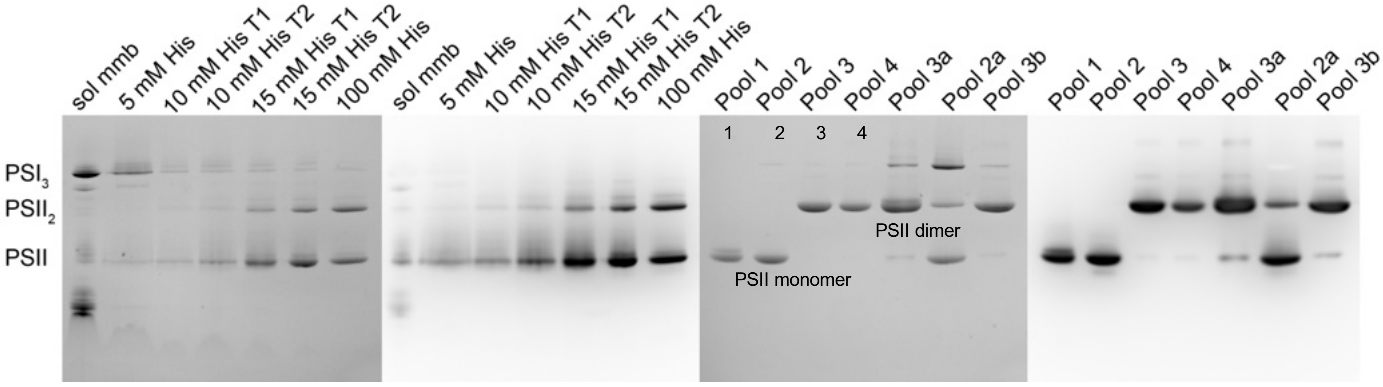


**Source data of Supplementary Fig 2 b, c**

a

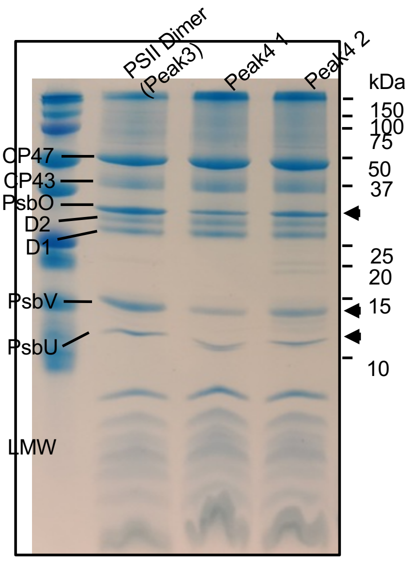


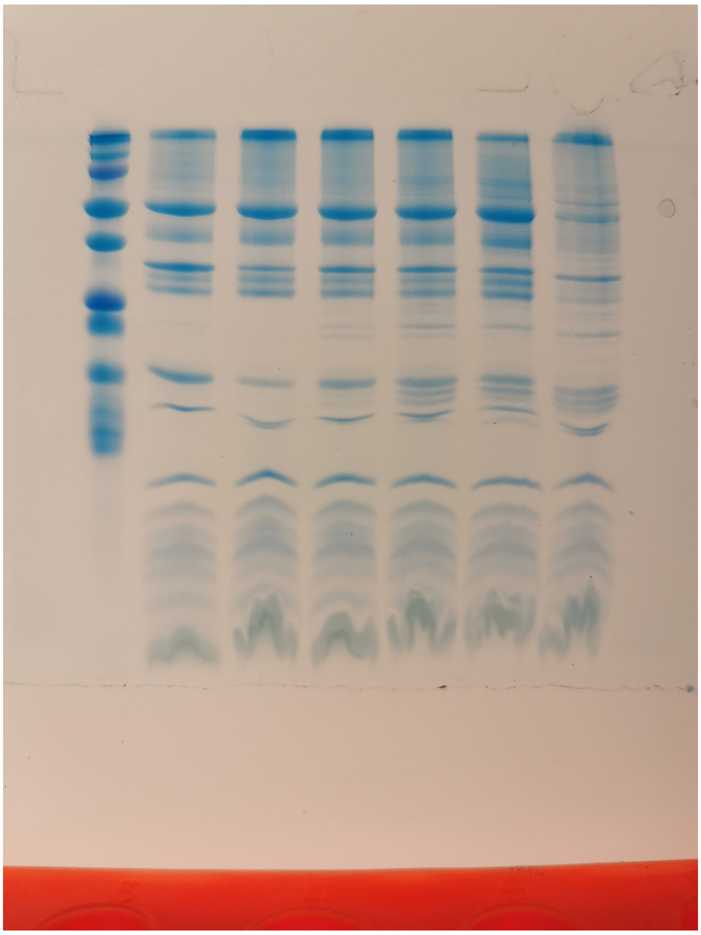

b

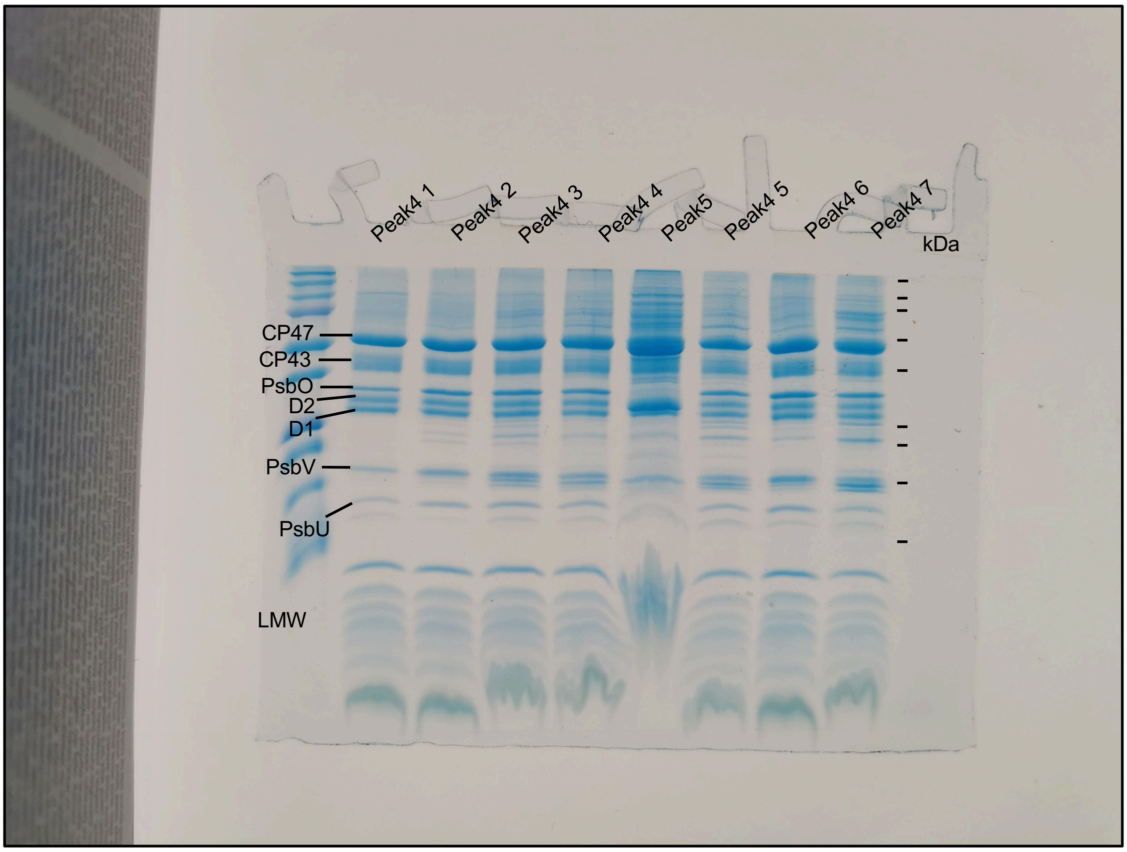


**Source data of Supplementary Fig 3 a, b**
